# Comprehensive computational modelling of the development of mammalian cortical connectivity underlying an architectonic type principle

**DOI:** 10.1101/426718

**Authors:** Sarah F. Beul, Alexandros Goulas, Claus C. Hilgetag

**Affiliations:** Institute of Computational Neuroscience, University Medical Center Hamburg-Eppendorf, Martinistr.52 - W36, 20246 Hamburg, Germany; Neural Systems Laboratory, Department of Health Sciences, Boston University, Commonwealth Ave. 635, 20115 Boston, MA, USA

## Abstract

The architectonic type principle attributes patterns of cortico-cortical connectivity to the relative architectonic differentiation of cortical regions. One mechanism through which the observed close relation between cortical architecture and connectivity may be established is the joint development of cortical areas and their connections in developmental time windows. Here, we describe a theoretical exploration of the possible mechanistic underpinnings of the architectonic type principle, by performing systematic computational simulations of cortical development. The main component of our *in silico* model was a developing two-dimensional cortical sheet, which was gradually populated by neurons that formed cortico-cortical connections. To assess different explanatory mechanisms, we varied the spatiotemporal trajectory of the simulated histogenesis. By keeping the rules governing axon outgrowth and connection formation constant across all variants of simulated development, we were able to create model variants which differed exclusively by the specifics of when and where neurons were generated. Thus, all differences in the resulting connectivity were due to the variations in spatiotemporal growth trajectories. Our results demonstrated that a prescribed targeting of interareal connection sites was not necessary for obtaining a realistic replication of experimentally observed connection patterns. Instead, we found that spatiotemporal interactions within the forming cortical sheet were sufficient if a small number of empirically well-grounded assumptions were met, namely planar, expansive growth of the cortical sheet around two points of origin as neurogenesis progressed, stronger architectonic differentiation of cortical areas for later neurogenetic time windows, and stochastic connection formation. Our study pinpointed potential mechanisms of how relative architectonic differentiation and cortical connectivity become linked during development. The successful prediction of connectivity in two species, cat and macaque, from simulated cortico-cortical connection networks further underscored the general applicability of mechanisms through which the architectonic type principle can explain cortical connectivity in terms of the relative architectonic differentiation of cortical regions.

**Author Summary:** The mechanisms that govern the establishment of cortico-cortical connections during the development of the mammalian brain are not completely understood. In computational simulation experiments reported here, we explored the foundations of an architectonic type principle, which attributes adult cortical connectivity to the relative architectonic differentiation of connected areas. Architectonic differentiation refers, among other characteristics, to the cellular make-up of cortical areas. This architectonic type principle has been found to account for diverse properties of cortical connectivity across mammalian species. Our *in silico* model generated connectivity patterns consistent with the architectonic type principle and typically observed in mammalian cortices, if model settings were chosen such that they corresponded to empirical observations about how cortical development proceeds. Our computational experiments systematically evaluated previously proposed mechanisms of cortical development and showed that connectivity consistent with the architectonic type principle arises only from realistic assumptions about the growth of the cortical sheet.

## Introduction

Axonal connections among cortical areas are the structural substrate of information transfer throughout the brain. These cortico-cortical connections form networks that are neither regular nor random, and exhibit large-scale topological features, such as modules and hubs [1, 2], rich-clubs [3, 4] and diverse-clubs [5], that have been the subject of wide-ranging investigations [6-19]. Moreover, there exist noteworthy regularities in the laminar patterns of cortical projection origins and terminations [20-23].

Many structural features of the cortex have been probed for their relationship to axonal connections between brain regions. For example, aspects of cell morphology have been shown to correlate with properties such as area degree (i.e., the number of projections maintained by an area) in the macaque monkey [24, 25] and humans [26].

### An architectonic type principle linking cortical structure and connectivity

One potent explanatory framework that imposes order onto the tangle of cortico-cortical connections is the so-called structural model [27]; reviewed in [28, 29], also termed architectonic type principle (ATP). This principle describes the patterns of cortical projections and their laminar origins and terminations in terms of the relative architectonic differentiation of brain areas. Briefly, graded differences in cortical architecture have been found to account for the graded patterns observed in the distribution of projection origins and targets across cortical layers [27, 30-36]. Moreover, greater similarity in the architectonic differentiation of cortical areas has been found to be associated with higher connection frequency between them, above and beyond the explanatory power of spatial proximity [34, 36, 37]; see [29, 38] for reviews). Originally described for ipsilateral connections of the macaque prefrontal cortex [27], the ATP has since been confirmed for a considerable number of brain systems and species, as well as contralateral connections [30-37, 39-42], suggesting a mammalian-general organisational principle. The general applicability of this principle was further supported in a recent study by performing prediction analyses that transferred information across mammalian species [43]. Specifically, by training a classifier on the relationship between cortical structure and connections in a first species, area-to-area connectivity in a second species could be reliably predicted from structural variations of cortical areas in the second species without making changes to the classifier.

The architectonic type principle, thus, allows the prediction of cortico-cortical connectivity from brain architecture regularities. Further substantiation of the ATP calls for a mechanistic explanation of how the described relationships between brain architecture and connectivity may emerge. From early on, the origin of this relationship has been hypothesized to be linked to developmental events [27]. Specifically, the observed close relationship between variations in cortical structure and axonal connections may arise from an interplay between the ontogenetic time course of neurogenesis and concurrent connection formation [29, 35, 44]. Areas which develop during different time windows were suggested to be afforded distinct opportunities to connect, with self-organisation rather than precisely targeted connection formation leading to the strikingly regular final connectivity patterns (also cf. [45]). Put differently, it has been hypothesized that spatiotemporal interactions in the forming tissue, and specifically the relative timing of neurogenesis across the cortex, determine the connectivity patterns between cortical areas. Empirically, such a relationship has, for example, been observed in the olfactory system of the rat [46].

Here, using systematic computational simulation experiments, we explored whether this suggested mechanism may be capable of generating cortico-cortical connectivity consistent with empirical observations and the architectonic type principle (Fig. 1). To this end, we implemented an *in silico* model of the growing two-dimensional cortical sheet of a single cerebral hemisphere that was progressively populated by neurons and divided into cortical areas. Model neurons randomly grew their axons across the cortical sheet and stochastically formed connections with potential postsynaptic targets (similar, for example, to simulation experiments in [47] and [48]). We assessed the resulting network of simulated structural connections between cortical areas in the same way as in previous experimental studies (e.g., [34, 36]) and compared the results to the empirical observations. Since we constrained the model to a single hemisphere, the simulated connections represent ipsilateral connectivity. Following this general approach, we characterized a number of variants of the *in silico* model of the growing cortical sheet, which differed in their adherence to empirical observations about developmental processes, specifically the spatiotemporal sequence of neurogenesis across the cortex. By comparing the networks generated from these variants, we could infer which aspects of the proposed mechanistic underpinnings of the ATP, particularly, which neurodevelopmental assumptions, were necessary to approximate empirical ipsilateral cortical connectivity.

**Fig 1.**
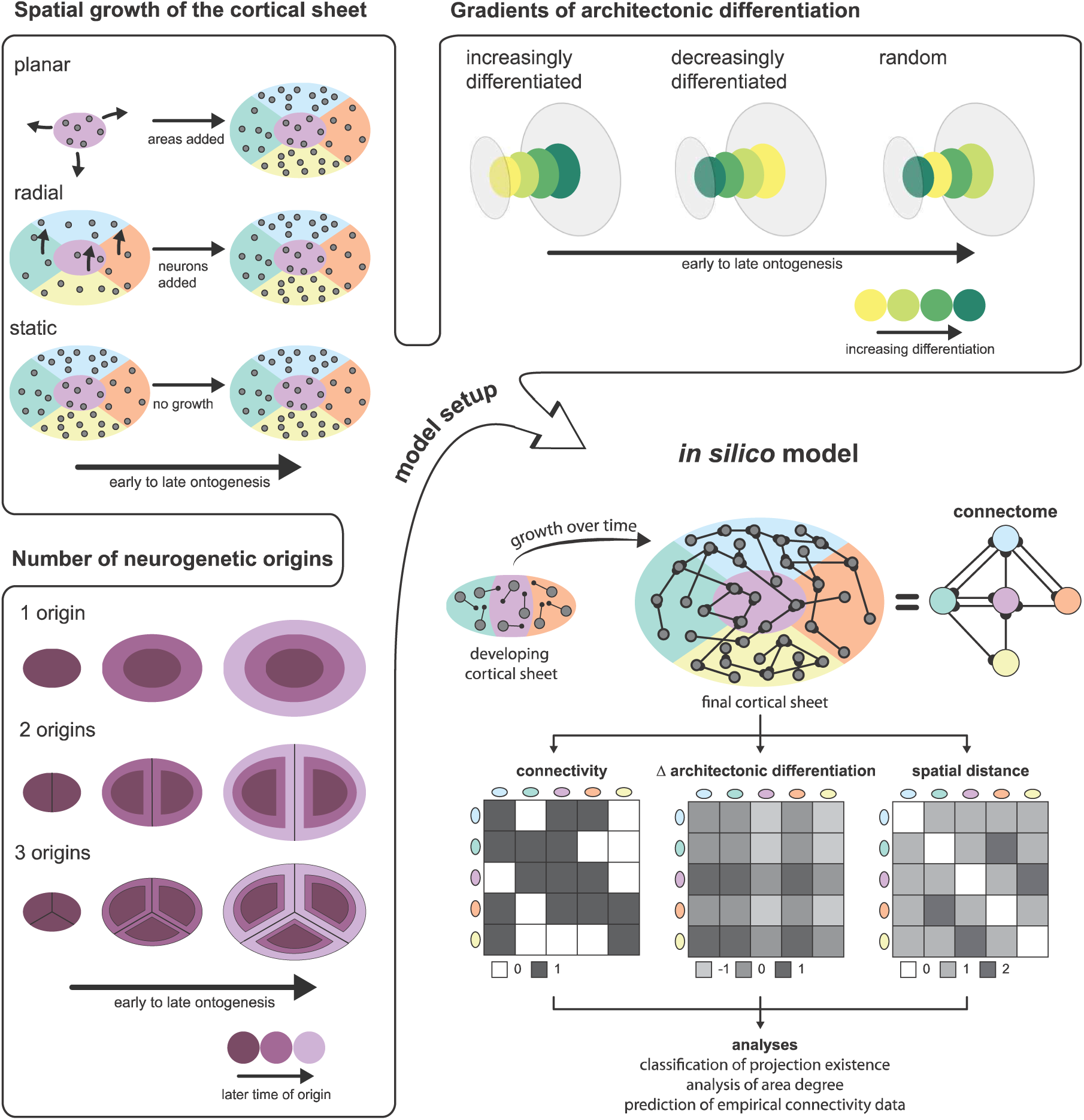
Neurodevelopmental assumptions and overview of the *in silico* model. The figure illustrates the assumptions regarding neurogenesis that were varied in the *in silico* model. The spatial growth of the cortical sheet of a single hemisphere was modelled in three possible ways: First, planar growth, in which the neurons comprising a cortical area develop at the same time and the cortical sheet expands as more areas materialize. Second, radial growth, in which neurons across the entire extent of the final cortical sheet develop at the same time, and the final complement of neurons is reached by gradual growth of neurons at a constant rate. Third, no growth, that is, a static cortical sheet on which the final complement of neurons is already present from the onset. Regarding the gradients of architectonic differentiation, we considered three possible relationships between the time at which an area was formed (time of neurogenesis) and its architectonic differentiation (approximated by neuron density). First, areas could be the more differentiated the later in ontogenesis they were formed (increasingly differentiated). This scenario corresponds to the *realistically oriented density gradient* we incorporated in the *in silico* model. Second, areas could be less differentiated, the later their time of origin was (decreasingly differentiated). This scenario corresponds to the *inversely oriented density gradient* in the *in silico* model. Third, there could be no gradient of differentiation aligned with neurogenetic timing, that is, the neuron density of newly formed areas varied randomly throughout ontogenesis. As a third factor that determined the spatiotemporal growth trajectory of the cortical sheet, we considered the number of neurogenetic origins. There could either be a single origin, such that more recently formed areas occupied the fringes of the cortical sheet, or there could be two or three origins. In this case, recently formed areas would be interleaved with areas that were formed earlier, as the neurogenetic origins were moved apart by addition of areas around them. From these assumptions on neurogenetic processes shaping the cortical sheet, we set up different variants of an *in silico* model in which axons grew randomly across the developing cortical sheet and stochastically formed connections. We translated the resulting neuron-level connectivity to area-level connectivity and extracted structural measurements from the simulated cortical sheet. As in previous studies of mammalian connectomes, we considered the difference in architectonic differentiation between areas and their spatial distance. Thus, we simulated sets of measures which we could then analyse in the same way as empirical data, and compared the results to empirical findings. Specifically, we used simulated architectonic differentiation and spatial distance to classify whether a connection existed in the final simulated network; we probed whether there was an association between simulated architectonic differentiation and the number of connections maintained by an area; and we used a classifier trained on the simulated data to predict connection existence in two sets of empirical connectivity data, from the cat and the macaque cortex.

### Aspects of neural development that prescribe spatiotemporal trajectories of cortical growth

We explicitly incorporated three aspects of corticogenesis in our simulations which are briefly described here.

First, the cortical sheet is established through neurogenesis spreading out from spatial origins, or primordial points, so that the surface of the cortex expands over time. This expansion is accompanied by a gradient in the time of onset of neurogenesis across the cortical sheet, which we refer to as the planar gradient of time of neurogenesis [49-59]. Developmental studies indicate that neurogenesis proceeds from at least two points of origin [56, 59, 60], with new neurons successively increasing the extent of cortical tissue between these neurogenetic origins. This progression entails that areas formed earlier become further separated on the cortical sheet as new areas are generated. Moreover, there is a superimposed radial gradient in the progression of neurogenesis [49, 50, 52, 61, 62], resulting in the characteristic inside-out generation sequence of neurons across layers (meaning that, with the exception of neurons in layer I, neurons in lower cortical layers are generated before neurons in upper cortical layers). In contradistinction to the findings outlining a planar gradient in the onset of neurogenesis, as described above, it has also been suggested that the onset of neurogenesis is simultaneous across the cortex [63, 64]. To contrast these two interpretations, we included both alternatives in our simulation experiments, as described in more detail below.

Second, cortical areas that are generated later are generally more architectonically differentiated [44, 59, 65, 66]; also briefly reviewed in [35]. Gradual changes in cortical architecture along two trends were described already several decades ago [67-71]; reviewed in [29, 38]. In brief, the two foci of least differentiated cortex are the allocortical, three-layered archicortex and paleocortex. These cortices are surrounded by periallocortex, where additional layers can be discerned, but without the clear laminar organisation found in the isocortex. Proisocortex, the next stage of differentiation, has a definite laminar organisation, but is missing a well-developed layer 4. Finally, there are different levels of isocortex with increasing demarcation of laminar boundaries and prominence of layer 4. More recently, changes in cell cycle kinetics across the forming cortical sheet and genetic correlates of the neurogenetic gradients have been described [57, 58, 72-74], which elucidate how gradual changes in cortical architecture are effected and provide an association between time of origin and architectonic differentiation. Particularly, a lengthening in the cell cycle along the planar neurogenetic gradient is accompanied by a successive increase in the proportion of progenitor cells differentiating into neurons with each cell cycle. In combination with the mentioned relation between time of origin and final laminar position of neurons, this mechanism results in a relatively increased number of supragranular layer neurons in later generated sections of the cortical sheet. Thus, a positive correlation can be observed between time of origin and neuron density across the cortex [66]. This link has been corroborated by findings in the human cortex, which directly traced systematic architectonic variation of the cortex to the timing of development [44]. A lengthening of the overall developmental time period, and with it the neurogenetic interval, appears to be responsible for increased neuron numbers both within the cortex of a given species, as well as across species which differ in their overall level of architectonic differentiation [65, 66, 75]. In fact, it has been suggested that cortical architecture correlates not only with neurogenetic time windows during ontogenesis, but also with the succession of architectural differentiation observed during brain evolution [59, 70]. This finding suggests that phylogenetic age has a bearing on architectural gradients. As mentioned above, it has repeatedly been reported that areas at similar points in the architectonic differentiation spectrum, as well as within the two described trends of architectonic progression, are preferentially linked, even if they are dispersed throughout the brain (also reviewed in [38]). The link to phylogeny, added to this correlation between architectonic progression and associated connectivity, thus, further points towards a developmental origin of the interrelations captured by the architectonic type principle.

The third aspect of neurogenesis which we incorporated into our simulations is that axon outgrowth starts concurrently with, or immediately after, neuronal migration [73, 76-79], and appears to be largely unspecific spatially [80]. We, therefore, assumed that connection formation starts as soon as neurons were generated. Further assumptions derived from these observations were that axons grow randomly across the cortical sheet (i.e., with no particular spatial orientation) and that they indiscriminately form connections once they are close enough to a potential target neuron, a mechanism that has been named Peters’s Rule [81, 82]. Thus, the process of connection formation can be described as stochastic, and has been simulated in this way in previous computational models of connection development (e.g., [48]). This mechanism entails that the probability of a neuron forming a connection is only dependent on the probability of its axon finding a target neuron. Since neurons that are far apart are separated by a larger number of neurons that could accommodate the axon, the probability of connecting to a target neuron is the lower, the larger the distance between two neurons. In effect, there is a positive correlation between the spatial proximity and connection probability of different neurons.

### An *in silico* model for assessing spatiotemporal growth trajectories

The spatiotemporal dynamics of corticogenesis that emerge from the combination of these empirically grounded assumptions were hypothesized to result in the establishment of realistic cortico-cortical connectivity. In particular, we expected interactions between the spatial and temporal aspects of neurogenesis to lead to the formation of connections which are consistent with the predictions of the architectonic type principle concerning the relationship between areas’ relative architectonic differentiation and connection frequencies. Our simulation experiments, thus, contribute the first systematic exploration of the neurodevelopmental mechanisms that have been hypothesized to underlie the ATP [27, 29, 35, 40].

In summary, we implemented several aspects of neurogenesis in an *in silico* model of the growing mammalian cerebral cortex. These aspects were then modified in some variants of the model, so that they either corresponded to, or violated, empirically observed phenomena. This strategy allowed us to compare the cortico-cortical connectivity resulting from hypothetical variants that differed in their assumptions, where some of these assumptions were empirically grounded and others were not. The approach enabled us to assess the merits of mechanisms which have been proposed to link cortical structure and connectivity through the ATP.

## Results

### Overview

We simulated the growth of cortico-cortical connections between areas of different neuron density according to a constant set of growth rules. We evaluated how closely the simulated connectivity corresponded to empirical observations made in mammalian connectomes when the physical substrate of the connections, that is, the simulated cortical sheet, developed along different spatiotemporal trajectories. To this end, we systematically varied the settings of our *in silico* model to construct a number of variants, which we refer to as spatiotemporal growth layouts. We considered five sets of growth layouts: (1: *realistically oriented density gradient*) planar growth of the cortical sheet, such that cortical areas were added around neurogenetic origins, with new areas having an increasingly higher neuron density (i.e., neuron density increased with distance from a point of origin); (2: *inverse density gradient*) planar growth of the cortical sheet, such that cortical areas were added around neurogenetic origins, but with new areas having increasingly lower neuron density (i.e., neuron density decreased with distance from a point of origin); (3: *radial)* no planar growth of cortical areas on the fringes of the cortical sheet, but gradual addition of neurons at a constant rate across the cortical sheet, which resulted in an ordered gradient of area neuron density that was the same as in sets 1 and 4; (4: *static*) no planar growth of cortical areas, but the same final gradient of area neuron density as in sets 1 and 3; (5: *random*) planar growth of the cortical sheet, such that cortical areas were added around neurogenetic origins, but with no ordered gradient of area neuron density, instead neuron density varied randomly between locations on the cortical sheet. For all five sets, we implemented three growth modes: (*1D 1row*) one-dimensional growth implemented with one row of areas; (*1D 2rows*) one-dimensional growth implemented with two rows of areas; and (*2D*) two-dimensional growth. For all five sets, all three growth modes were implemented with planar growth around two neurogenetic origins. For set 1 (*realistically oriented density gradient*), we additionally implemented each growth mode with one neurogenetic origin as well as three (1D growth) or four (2D growth) neurogenetic origins. Thus, in total, we considered 21 growth layouts, grouped into five sets according to the spatiotemporal trajectory the cortical sheet traversed (see Fig. 2 and Table 1 for an overview).

We first present some general statistics of the simulated connectivity and then go on to characterize how well the relationship between connectivity and the two factors of (relative) neuron density and spatial distance corresponded to previously published empirical observations for the different growth layouts. Finally, we assess how well the different growth layouts predicted empirical connectivity, as an indication of how realistic the simulated connectivity was for a given growth layout. Figure 3 provides an outline of this procedure. Table 2 gives an overview of all results.

**Fig 2.**
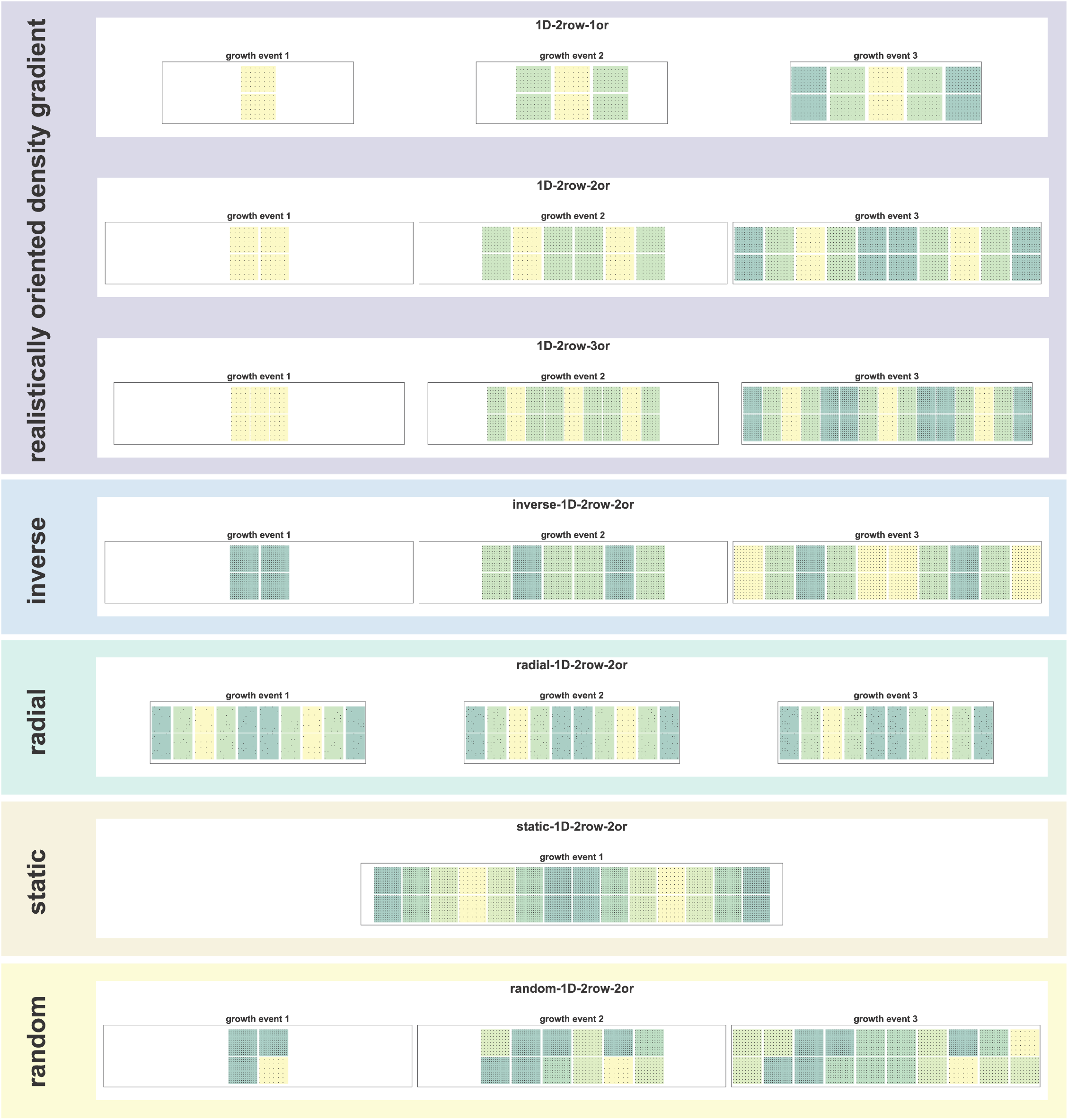
Developmental trajectories of growth layouts. The figure illustrates the spatiotemporal growth trajectory for different growth layouts. The successive population of the cortical sheet with neurons is shown for the first three growth events. For static growth, all neurons grow simultaneously, hence only one growth event is shown. Here, all growth layouts of growth mode *1D 2 rows* are shown. See Supplementary Figure S1 for an illustration of the developmental trajectories of all 21 growth layouts.

**Fig 3.**
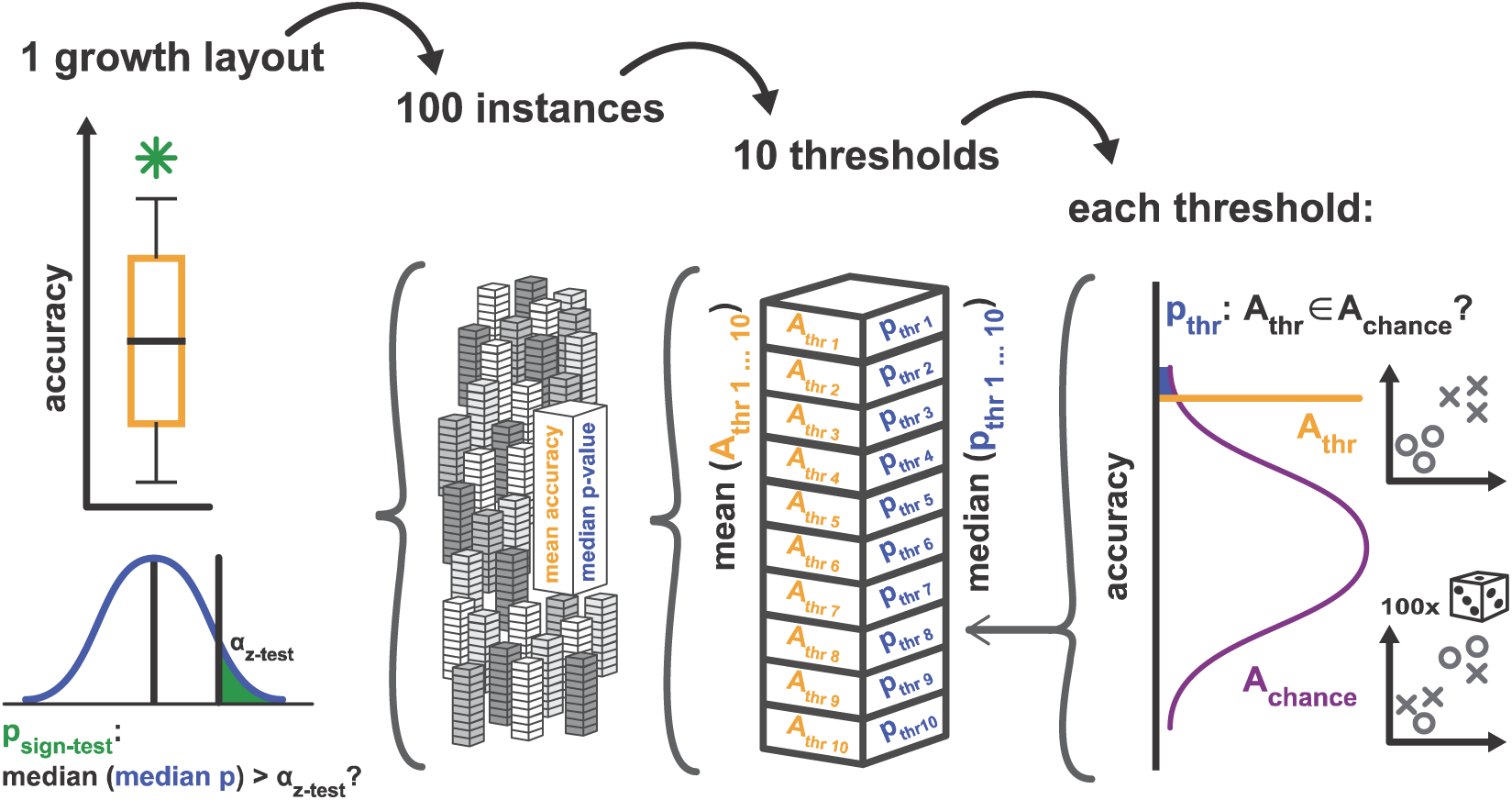
Validation procedure for measures of simulation-to-empirical classification performance. The figure illustrates the general procedure for assessing the performance of the classification of empirical data from the cat and macaque cortex by classifiers that were trained on simulated data; see main text for details. We computed median measures of classification performance for each growth layout and compared these measures against chance performance, as assessed by a permutation analysis. Specifically, for each of the 21 growth layouts shown in Figure 8 and Table 4, 100 instances were simulated. For each instance, classification was performed using 10 different classification thresholds. For each threshold, a simulation-trained classifier assigned labels to the empirical data, resulting in A_thr_. Additionally, a distribution of chance performance accuracies, A_chance_, was generated by classifying 100 times from randomly permuted non-sensical labels. A z-test quantified the probability that A_thr_ was an element of the distribution of A_chance_. The corresponding p-value p_thr_ was used for further calculations. For each simulation instance, classification performance from all 10 thresholds was averaged, resulting in one mean accuracy value and one median value of p_thr_ per instance, thus amounting to a total of 100 values each per growth layout. Figure 8 shows the distribution of mean accuracy values from these 100 instances, and indicates the median accuracy. The indication of significance in Figure 8 refers to the p-value obtained from a sign-test which assessed whether the median of the distribution of median values of p_thr_ was larger than the chosen significance threshold α_z-test_ of 0.05 (with a small value of p_sign-test_ indicating that p_thr_ was very unlikely to be larger than α_z-test_). Table 4 includes the median accuracy, median z-test p-value and the result of the sign-test. Shown here for accuracy, the procedure was analogous for the Youden index *J*, which is shown in Figure 9 and Table 4.

**Table 1.** Growth layouts. This table indicates the set, growth mode and number of neurogenetic origins for each of the 21 growth layouts. For each set, the determining properties of the spatiotemporal growth trajectory are indicated. Moreover, for each growth layout the total numbers of areas, growth events and neurons are included. Abbreviations and background colours introduced here are used throughout the figures.

| set | growth mode | # origins | final gradient of<br>neuron density<br>around origins | planar<br>growth of<br>cortical<br>sheet | radial<br>growth of<br>cortical<br>sheet | number<br>of areas | number of<br>growth<br>events | total<br>number of<br>neurons | abbreviation |
| --- | --- | --- | --- | --- | --- | --- | --- | --- | --- |
| realistically<br>oriented<br>gradient | 1D 1 row | 1 | realistically<br>oriented | ✓ | ✗ | 25 | 12 | 24897 | 1D-1row-1or |
|  | 1D 2 rows | 1 |  |  |  | 50 | 12 | 49794 | 1D-2row-1or |
|  | 2D | 1 |  |  |  | 81 | 5 | 40838 | 2D-1or |
|  | 1D 1 row | 2 |  |  |  | 26 | 6 | 26550 | 1D-1row-2or |
|  | 1D 2 rows | 2 |  |  |  | 52 | 6 | 53100 | 1D-2row-2or |
|  | 2D | 2 |  |  |  | 162 | 5 | 81676 | 2D-2or |
|  | 1D 1 row | 3 |  |  |  | 27 | 4 | 28215 | 1D-1row-3or |
|  | 1D 2 rows | 3 |  |  |  | 54 | 4 | 56430 | 1D-2row-3or |
|  | 2D | 3 |  |  |  | 196 | 4 | 100248 | 2D-4or |
| inverse<br>gradient | 1D 1 row | 2 | inverse | ✓ | ✗ | 26 | 6 | 23910 | inverse-1D-1row-2or |
|  | 1D 2 rows | 2 |  |  |  | 52 | 6 | 47820 | inverse-1D-2row-2or |
|  | 2D | 2 |  |  |  | 162 | 5 | 38994 | inverse-2D-2or |
| radial | 1D 1 row | 2 | realistically<br>oriented | ✗ | ✓ | 26 | 6 | 26550 | radial-1D-1row-2or |
|  | 1D 2 rows | 2 |  |  |  | 52 | 6 | 53100 | radial-1D-2row-2or |
|  | 2D | 2 |  |  |  | 162 | 5 | 81676 | radial-2D-2or |
| static | 1D 1 row | 2 | realistically<br>oriented | ✗ | ✗ | 26 | 1 | 26550 | static-1D-1row-2or |
|  | 1D 2 rows | 2 |  |  |  | 52 | 1 | 53100 | static-1D-2row-2or |
|  | 2D | 2 |  |  |  | 162 | 1 | 81676 | static-2D-2or |
| random | 1D 1 row | 2 | no gradient /<br>random | ✓ | ✗ | 26 | 6 | 26550 | random-1D-1row-2or |
|  | 1D 2 rows | 2 |  |  |  | 52 | 6 | 53100 | random-1D-2row-2or |
|  | 2D | 2 |  |  |  | 162 | 5 | 81676 | random-2D-2or |

**Table 2.** Summary of correspondence between simulation results and empirical observations. This table provides an estimate of the extent to which the connectivity resulting from each growth layout corresponds to expectations derived from empirically observed phenomena. ✓: good correspondence, ?: inconclusive, ×: correspondence not satisfactory.

| set | growth mode | # origins | connectivity between<br>areas of similar | classification of<br>connections: | number of<br>connections | classification of connections:<br>empirical from simulation |
| --- | --- | --- | --- | --- | --- | --- |

|  |  |  | neuron density | simulation from simulation |  | accuracy | Youden index *J* |
| --- | --- | --- | --- | --- | --- | --- | --- |
| realistically oriented gradient | 1D 1 row | 1 | ✓ | ✓ | ✓ | ? | ✓ |
|  | 1D 2 rows | 1 | ✓ | ✓ | ✓ | ? | ✓ |
|  | 2D | 1 | ✓ | ✓ | ✗ | ✓ | ✗ |
|  | 1D 1 row | 2 | ✓ | ✓ | ✓ | ✓ | ✓ |
|  | 1D 2 rows | 2 | ✓ | ✓ | ✓ | ✓ | ✓ |
|  | 2D | 2 | ✓ | ✓ | ✗ | ✓ | ✓ |
|  | 1D 1 row | 3 | ✓ | ✓ | ✓ | ✓ | ✓ |
|  | 1D 2 rows | 3 | ✓ | ✓ | ✓ | ✓ | ✓ |
|  | 2D | 3 | ✓ | ✓ | ✗ | ✓ | ✓ |
| inverse gradient | 1D 1 row | 2 | ✓ | ? | ✗ | ? | ? |
|  | 1D 2 rows | 2 | ✓ | ? | ✗ | ✗ | ✗ |
|  | 2D | 2 | ✓ | ? | ✗ | ? | ✗ |
| radial | 1D 1 row | 2 | ✗ | ✗ | ✗ | ? | ✗ |
|  | 1D 2 rows | 2 | ? | ✗ | ✗ | ? | ✗ |
|  | 2D | 2 | ✓ | ✗ | ✗ | ✗ | ✗ |
| static | 1D 1 row | 2 | ✗ | ✗ | ✗ | ✗ | ✗ |
|  | 1D 2 rows | 2 | ✗ | ✗ | ✗ | ✗ | ✗ |
|  | 2D | 2 | ✓ | ✗ | ✗ | ✓ | ✗ |
| random | 1D 1 row | 2 | ✗ | ✗ | ✗ | ? | ✗ |
|  | 1D 2 rows | 2 | ✗ | ✗ | ✗ | ? | ✗ |
|  | 2D | 2 | ✓ | ✗ | ✗ | ✗ | ✗ |
| details in |  |  | Fig. 5 | Fig. 6 | Fig. 7 | Fig. 8 | Fig. 9 |
| corresponding measure |  |  | correlation of relative connection frequency vs \|density difference\| | McFadden's Pseudo R<sup>2</sup> for \|density difference\| | correlation of area degree vs density | classification of connections in cat and macaque cortex: accuracy | classification of connections in cat and macaque cortex: Youden index *J* |

### Connection statistics

The cortico-cortical networks resulting from the simulations showed realistic levels of overall connectivity, with between 39% and 66% of possible connections present (Fig. 4A, Table 3). Previously, between 50% and 77% of connections were reported to be present in the macaque and cat cortex [22, 34, 83]. Some *2D* growth layouts reached higher levels of connectivity, with up to 87% of possible connections present. This connection density translated into several hundreds of inter-areal connections (Fig. 4B, Table 3), with between 250 and 400 connections for growth mode *1D 1row* and between 900 and 1500 connections for growth mode *1D 2rows*. Due to the large number of areas, connection numbers were much higher for *2D* growth layouts, between 8000 and 18600.

**Fig 4.**
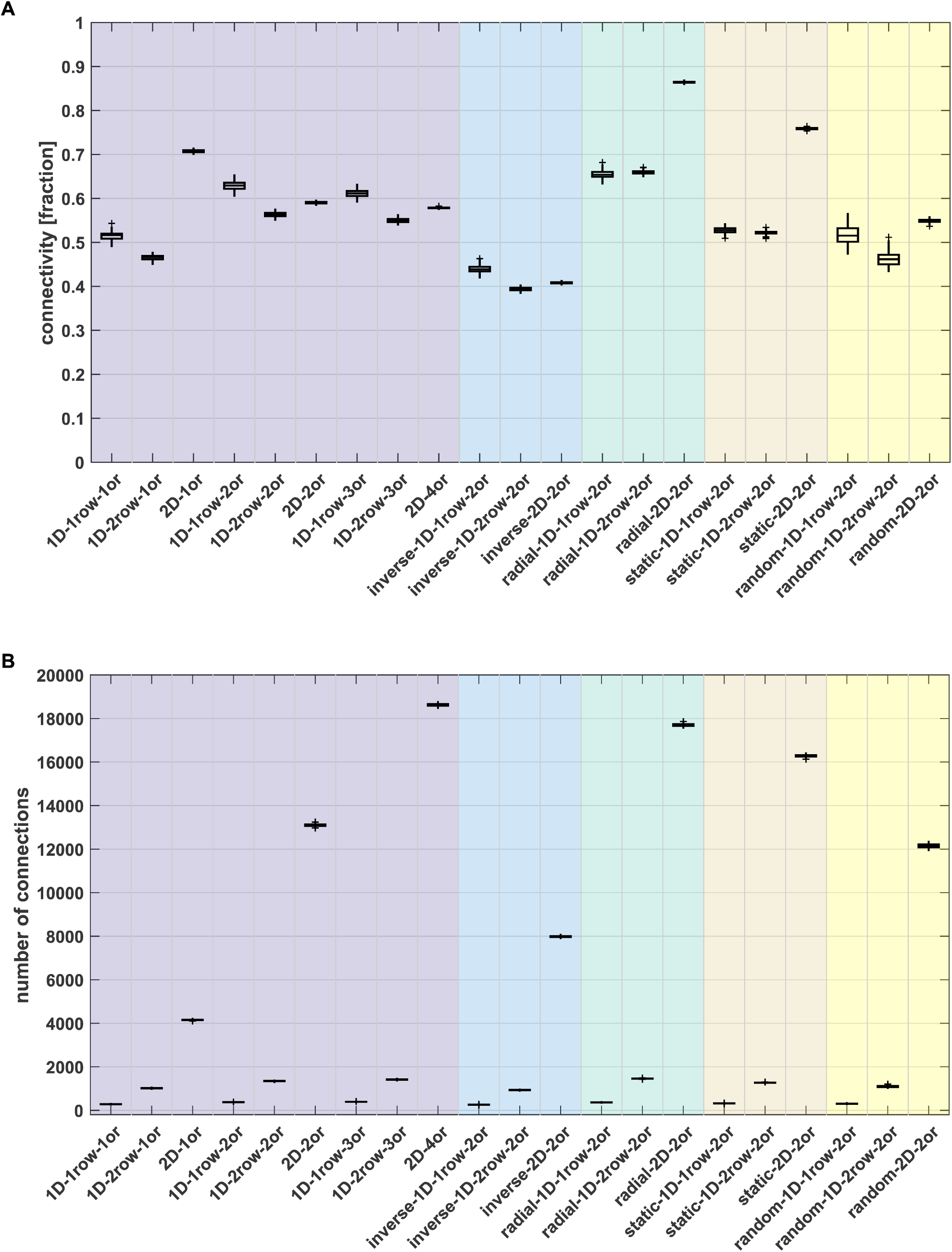
Connection statistics. (A) Percentage of connected areas, plotted as the fraction of possible connections that are present in the final simulated network. (B) Total number of connections among all areas. Box plots show distribution across 100 simulation runs per growth layout, indicating median (line), interquartile range (box), data range (whiskers) and outliers (crosses, outside of 2.7 standard deviations). See Table 3 for a summary. Abbreviations and background colours as in Table 1.

**Table 3.**
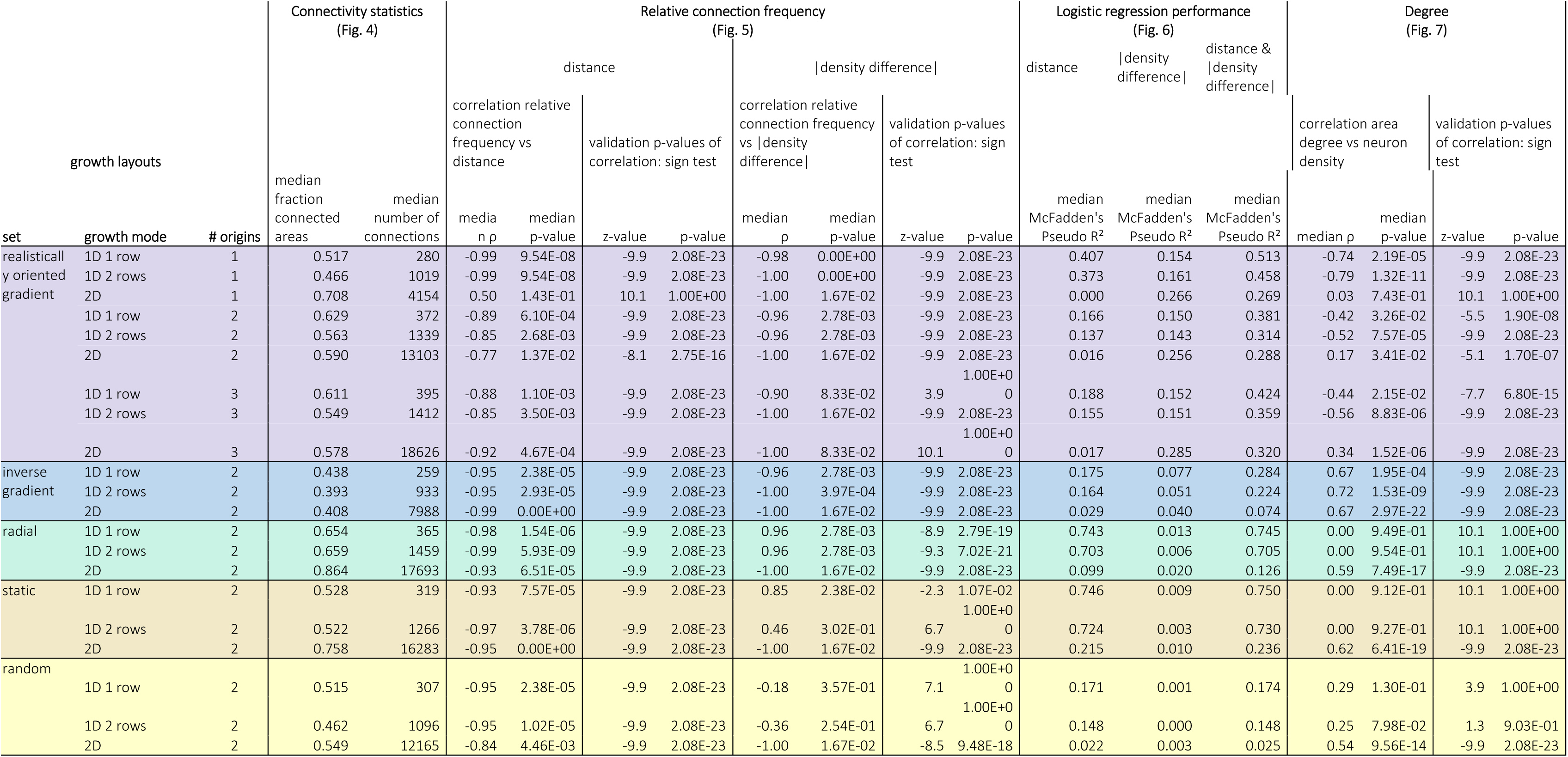
Summary connectivity statistics, correlation with relative projection frequency, classification performance logistic regression, and correlation with area degree. This table lists the median values indicated by the box plots in Figures 4 to 7. Where applicable, the table additionally lists the associated median p-value of Spearman rank correlations as well as the z-value and p-value of a left-tailed sign test testing the distribution of rank correlation p-values for a median of α = 0.05. Background colours as in Table 1.

### Contributions of distance and density difference to connectivity patterns

We first checked how well the simulated networks corresponded to the empirical observations that a larger fraction of connections is present between regions that are more similar in neuronal density, as suggested by the architectonic type principle, and spatially closer to each other. To this end, we computed the relative frequency of present connections (Fig. 5, Table 3). We then examined how well both factors, absolute density difference and distance, enabled the reconstruction of the simulated networks using logistic regression. Specifically, we assessed this by computing McFadden’s Pseudo R^2^ statistic, which provides a measure of the increase in the model log-likelihood with inclusion of either or both factors compared to a null model (Fig. 6, Table 3).

**Fig 5.**
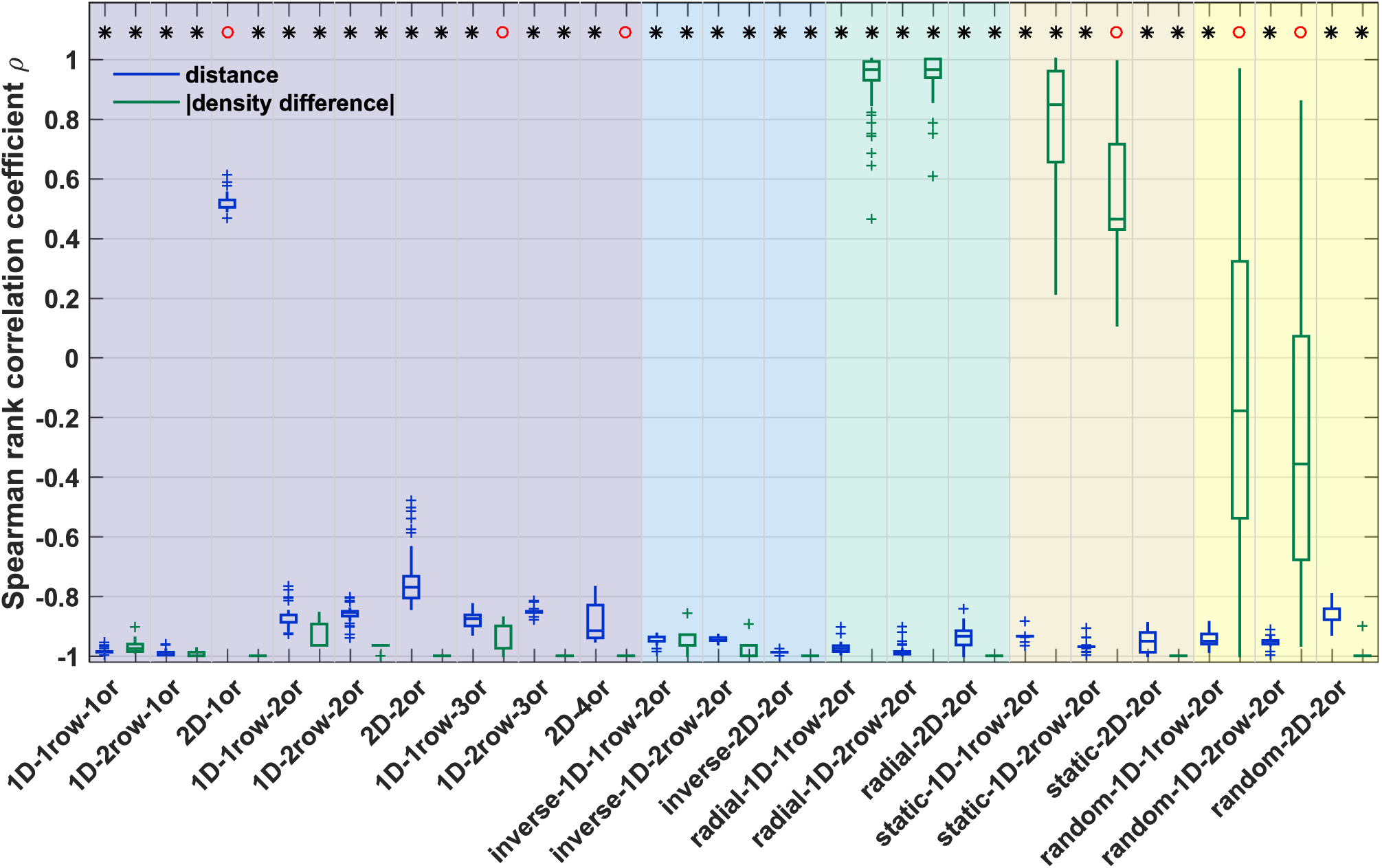
Correlation of distance and absolute density difference with relative connection frequency. Spearman rank correlation coefficients are provided for the correlation between relative connection frequency and distance (blue) or absolute density difference (green). A sign test was used to test whether the distribution of associated Spearman rank correlation p-values had a median value smaller than α = 0.05. The result of the sign test is indicated on top; black star: median p < 0.05, red circle: median p >= 0.05. See Supplementary Figure S2 for representative plots of the correlation for individual simulation runs. Box plots show distribution across 100 simulation runs per growth layout, indicating median (line), interquartile range (box), data range (whiskers) and outliers (crosses, outside of 2.7 standard deviations). See Table 3 for a summary. Abbreviations and background colours as in Table 1.

**Fig 6.**
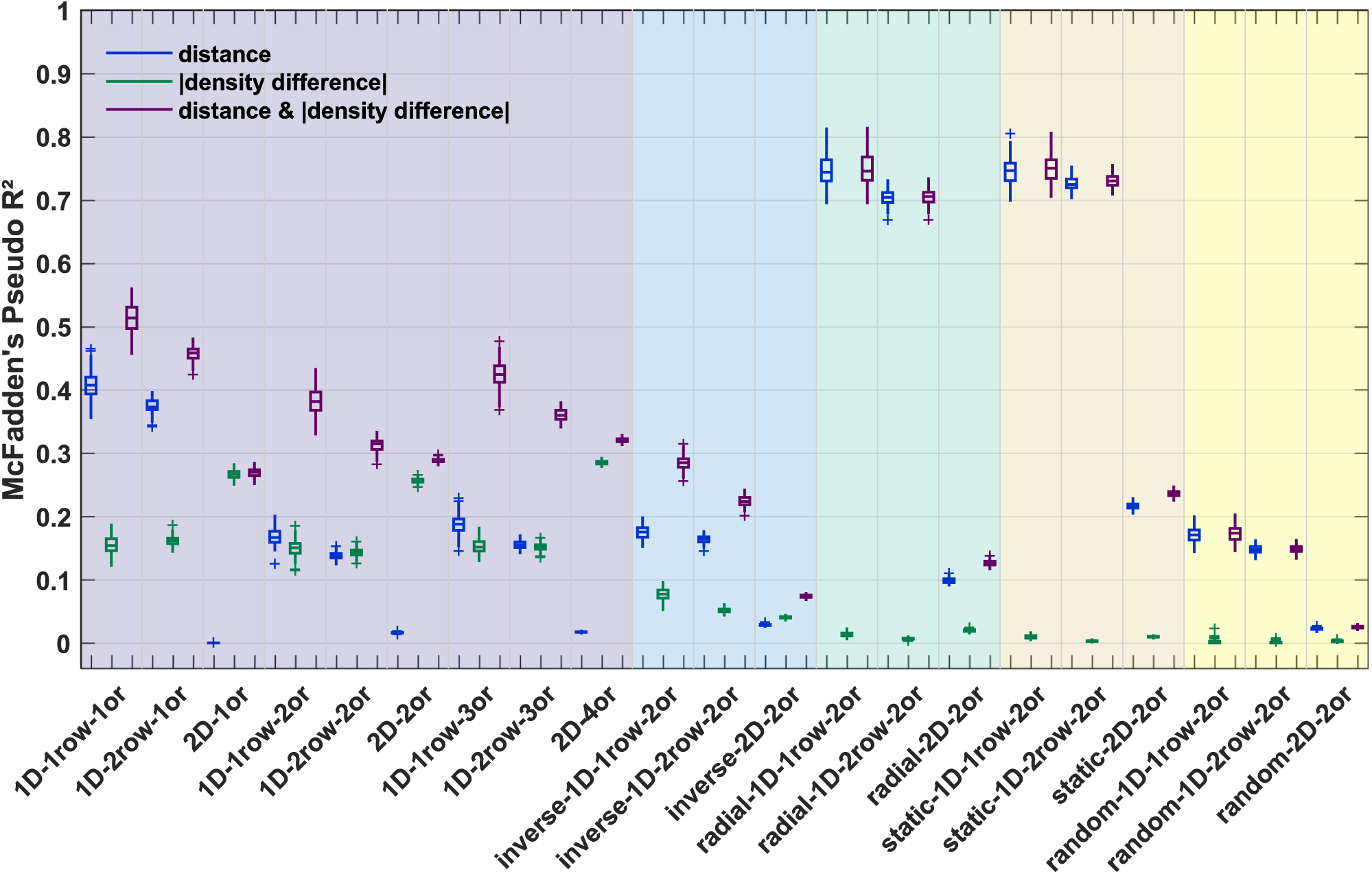
Logistic regression performance for classification of simulation data from simulation data. Within each growth layout, a logistic regression was performed to classify connection existence from three sets of factors: distance (blue), absolute density difference (green), or distance as well as absolute density difference simultaneously (purple). To assess whether classification performance was better than chance, McFadden’s Pseudo R^2^ was computed against performance of a null-model, where a constant was the only factor included in the logistic regression. Box plots show distribution across 100 simulation runs per growth layout, indicating median (line), interquartile range (box), data range (whiskers) and outliers (crosses, outside of 2.7 standard deviations). See Table 3 for a summary. Abbreviations and background colours as in Table 1.

#### Relative frequency of present connections

In general, connections were more likely to be present across smaller distances (Fig. 5, Supplementary Fig. S2). The relative frequency of present connections was very strongly negatively correlated with the distance between areas. The correlation was significant for all growth layouts, except for the *2D 1origin* growth layout. This effect was due to very weak connections being formed across even the longest distances in this growth layout, which resulted in a moderate positive correlation that did not reach significance. However, also for this growth layout, the correlation became strongly negative and significant if connections with fewer than 10 constituent axons were excluded, in line with previous treatment of empirical data [31, 83].

In contrast, the correlation of relative connection frequency with density difference was not uniform across all growth layouts. For *1D random, static* and *radial* growth layouts, the density difference was not significantly or else positively correlated with relative connection frequency. For *2D* growth layouts, however, the correlation was negative and significant for all three of those sets.

Conversely, the density difference was very strongly negatively correlated with relative projection frequency for all growth layouts with oriented growth (i.e., *realistically oriented gradient* and *inverse gradient*). The only exceptions here were the *1D 1row 3origins* growth layout and the *2D 4origins* growth layout. For reasons of computational efficiency, these layouts were implemented with only five and four density difference tiers, respectively. For the *1D 1row 3origins* growth layout, the deviation of relative connection frequency from a perfect negative correlation in one of the five tiers was, therefore, sufficient to render the rank correlation insignificant, with a p-value of 0.083. Similarly, for the *2D 4origins* growth layout, the minimal p-value that could be obtained from a rank correlation across the four tiers was 0.083, which is not low enough to reach significance. However, the correlation coefficients for both growth layouts consistently indicated a very strong to perfect negative correlation (cf. also Supplementary Fig. S2).

### Logistic regression

When we predicted connection existence using binary logistic regression, the inclusion of distance as a predicting factor markedly increased prediction performance as compared to the constant-only null model, with median McFadden’s Pseudo R^2^ values of at least 0.14 (Fig. 6). This was not true for the *2D* growth layouts with planar growth of the cortical sheet (i.e., the *static* and *radial 2D* growth layouts are excepted here), where distance did not markedly increase prediction performance compared to the constant-only null model, with median McFadden’s Pseudo R^2^ values of at most 0.03. For the *radial 2D* growth layout, distance performed intermediately with a median McFadden’s Pseudo R^2^ value of 0.10, indicating moderate performance. Absolute density difference as the only predictive factor did not increase prediction performance compared to the constant-only null model for all *random, static* and *radial* growth layouts, with median McFadden’s Pseudo R^2^ values below 0.03. However, inclusion of absolute density difference led to an increase in prediction performance for the growth layouts with oriented growth. For the growth layouts with a *realistically oriented density gradient*, the performance increase was moderate to very high, with median McFadden’s Pseudo R^2^ values between 0.14 and 0.28. For growth layouts with an *inverse density gradient*, in contrast, the performance increase was very small, with median McFadden’s Pseudo R^2^ values between 0.04 and 0.08.

Including distance and absolute density difference jointly as predictors for the logistic regression led to a moderate to very high increase in prediction performance compared to the constant-only null model, with median McFadden’s Pseudo R^2^ values of at least 0.13, but mostly above 0.20 and up to 0.75. The only exceptions to this finding were the *random* and the *inverse 2D* growth layouts, which did not reach median McFadden’s Pseudo R^2^ of 0.10.

In summary, a binary logistic regression adequately allowed to predict connection existence from distance and absolute density difference for the overwhelming majority of growth layouts. This result was to be expected given the rules of connection growth that governed the formation of the simulated networks. The notable dissociation that could be observed in the separate prediction from distance and density difference was that distance markedly contributed to prediction performance for most growth layouts, while the contribution of density difference was more specific. Namely, density difference most strongly allowed prediction of connection existence for the layouts with oriented growth of the cortical sheet and a *realistically oriented density gradient*.

### Number of connections per area

Another property of the simulated networks that we compared to empirical observations was area degree (i.e., the number of connections per area). We previously reported that, in biological cortical networks, the number of connections maintained by an area is negatively correlated with the area’s cytoarchitectonic differentiation [34, 36]. Here, we show an analogous analysis for the simulated networks (Fig. 7, Table 3, Supplementary Fig. S3). For *random*, *static* and *radial* growth layouts, area degree was not significantly correlated with neuron density, with the exception of *2D* growth layouts, which showed a positive and significant correlation in each case.

**Fig 7.**
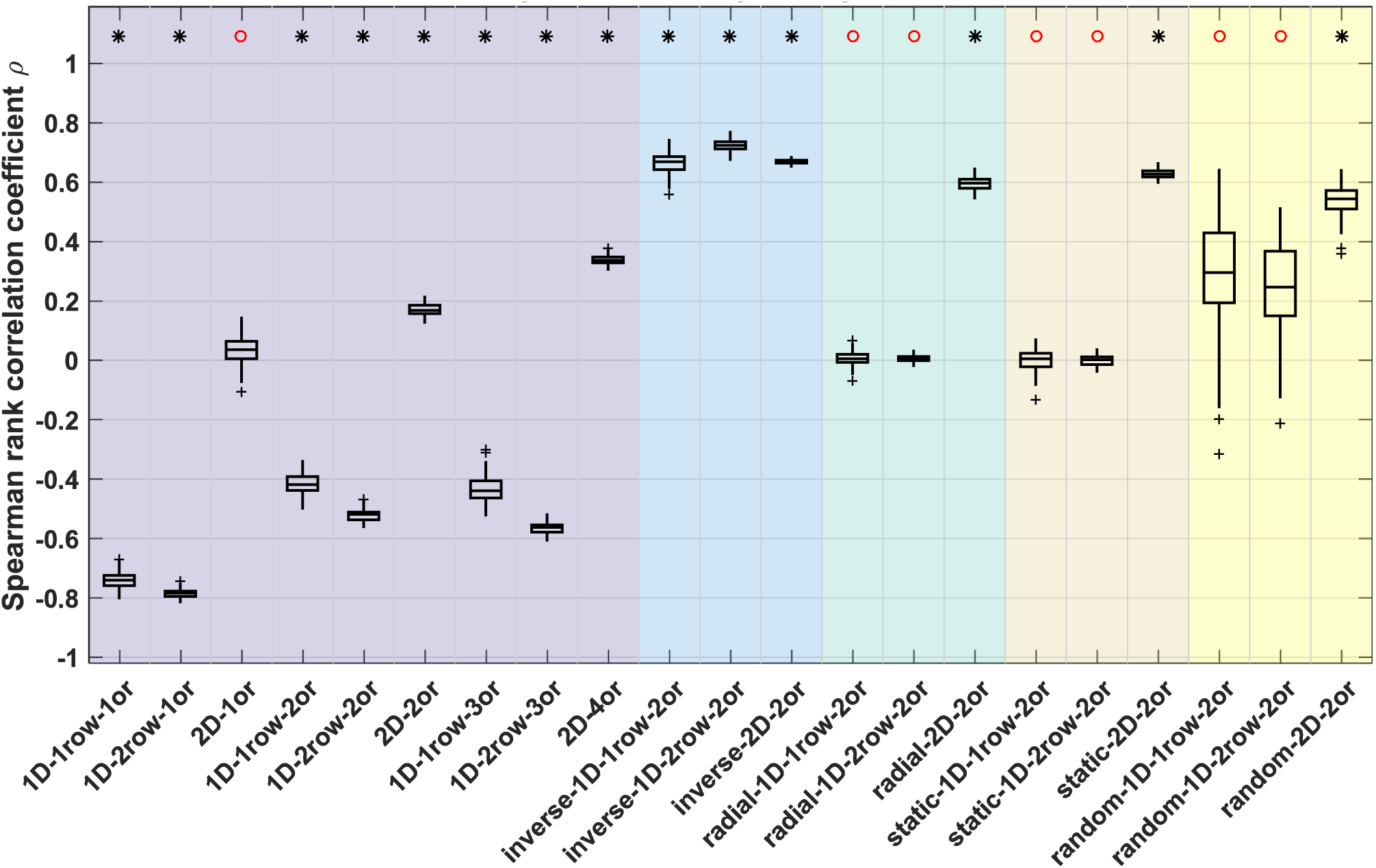
Correlation of area degree with neuron density. Spearman rank correlation coefficients for the correlation between area degree (number of connections) and area neuron density. A sign test was used to test whether the distribution of associated Spearman rank correlation p-values had a median value smaller than α = 0.05. The result of the sign test is indicated on top; black star: median p < 0.05, red circle: median p >= 0.05. See Supplementary Figure S3 for representative plots of the correlation for individual simulation runs. Box plots show distribution across 100 simulation runs per growth layout, indicating median (line), interquartile range (box), data range (whiskers) and outliers (crosses, outside of 2.7 standard deviations). See Table 3 for a summary. Abbreviations and background colours as in Table 1.

Growth layouts with *realistically oriented density gradients* showed a strongly negative, significant correlation between area degree and neuron density, with median correlation coefficients between −0.42 and −0.79 for both *1D* growth modes. Conversely, for growth layouts with an *inverse density gradient*, area degree was strongly positively correlated with neuron density. For *2D* growth along a *realistically oriented density gradient*, the observed effect was more variable. Correlation coefficients were of weak to moderate magnitude, and the correlation was not significant for *2D* growth around one origin (*2D 1origin:* median ρ = 0.03, median p > 0.05; *2D 2origins:* median ρ = 0.17, median p < 0.05; *2D 4origins:* median ρ = 0.34, median p < 0.05). This observation was in contrast to the strongly positive and significant correlations observed for the *2D* growth layouts without oriented growth, where median correlation coefficients were larger than 0.50 (*random 2D:* median ρ = 0.54; *static 2D:* median ρ = 0.62; *radial 2D:* median ρ = 0.59). We, therefore, concluded that the effect of oriented growth along a *realistically oriented density gradient* on area degree, as observed for both *1D* growth modes, persisted in the *2D* growth mode, but that it was not strong enough to completely abolish the tendency for a positive correlation between area degree and neuron density inherent to the *2D* growth layouts, instead only diminishing the positive correlation.

In summary, the empirically observed negative correlation between area degree and neuron density was only reproduced for the growth layouts with a *realistically oriented density gradient*. We cannot rule out that there existed a minor contribution of geometric centrality to this relationship. However, taking into account the results for the *radial* and *static* growth layouts made clear that such an effect, if there was any in the *realistically oriented gradient* growth layouts, could only be secondary. Without expansive, planar growth, there is no temporal advantage helping earlier-formed areas to accrue more connections. Any negative correlation between neuron density and area degree would, thus, be caused by geometric centrality. Figure 7 illustrates that no such correlation arises, instead area degree appears to vary randomly with neuron density for the *radial* and *static* growth layouts.

### Prediction of empirical connectivity data from simulated networks

In the previous sections, we showed that empirically observed regularities, particularly a close relationship between connection existence and the two factors of relative neuron density and spatial distance, could be reproduced *in silico*. We further characterized how well the simulation captured this phenomenon by predicting empirical connectivity using classifiers trained on the simulated networks. Classification performance was used as a measure of how well the properties of the artificially generated networks reflected the characteristics of empirical brain networks, in particular, the macaque and cat cortical connectomes. We report two measures of classification performance, accuracy and the Youden index, *J*. Accuracy was calculated as the percentage of predictions that were correct, while the Youden index is a summary measure that takes into account both the rate of true positives and the rate of true negatives and indicates divergence from chance performance.

As seen from Figures 8 and 9, classification performance was generally higher for the macaque connectome than for the cat connectome. However, the described differences between growth layouts held for both species. We also provide the fraction of the available empirical connections that were classified in each species (Fig. 10, Table 4). Generally, between 30% and 60% of the empirical connections were classified, with some growth layouts reaching up to 86% (Fig. 10). However, for some growth layouts, nearly no empirical connections reached posterior probabilities of at least 0.75 (the minimal threshold applied for assigning a predicted label), and, thus, very low fractions of the available empirical connections were classified. Specifically, this applied to *random* growth layouts (median fraction classified between <0.01 and 0.14) and the *inverse 2D* growth layout (median fraction classified macaque: 0.08, cat: 0.05). The overall low posterior probabilities for these growth layouts and the resulting small fraction of classified empirical connections already suggested that the properties of those layouts did not correspond well to the properties of the empirical networks. This impression was corroborated by other measures (see below).

**Fig 8.**
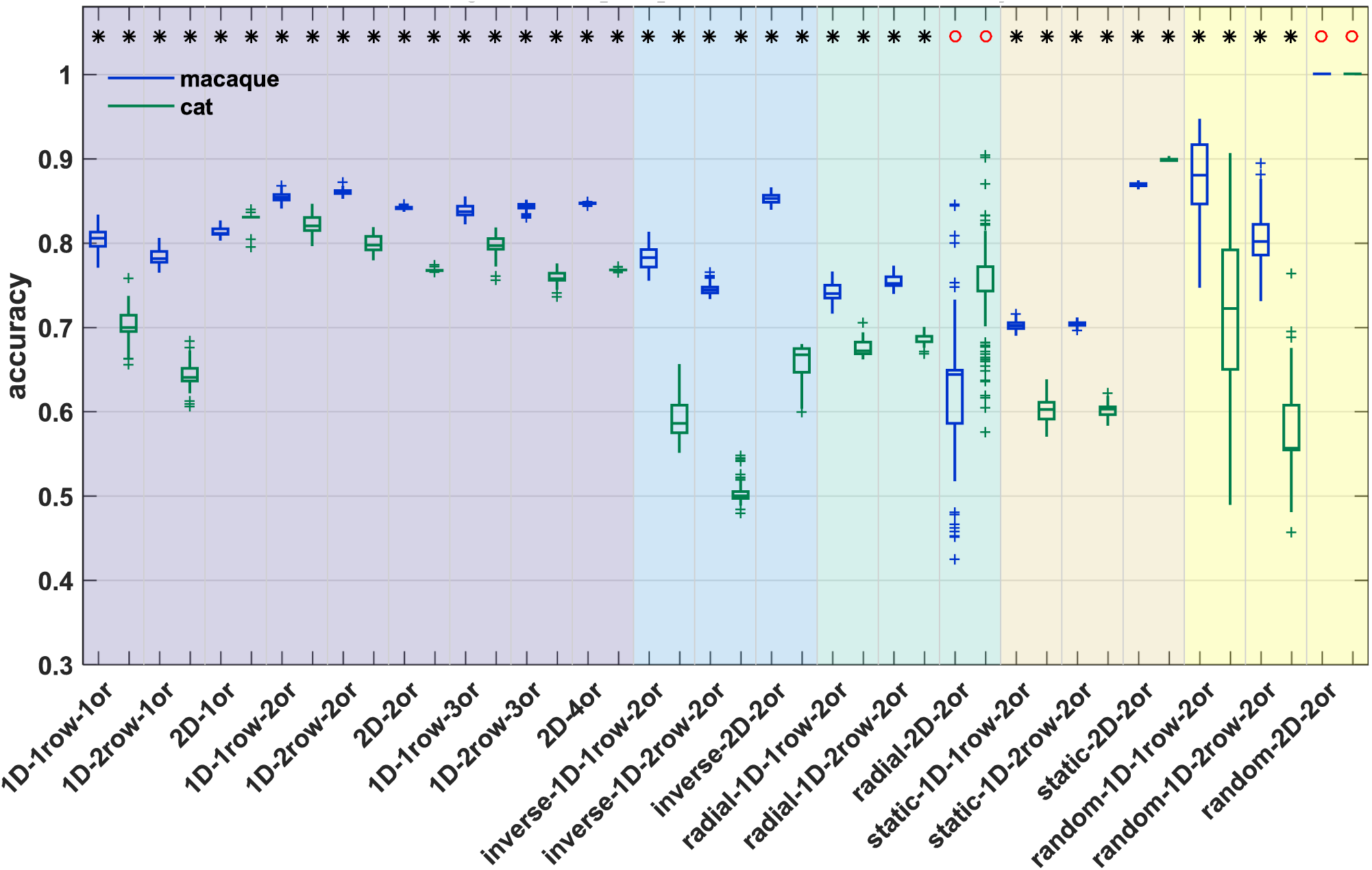
Classification accuracy for prediction of empirical connection existence from simulation data. A classifier was trained to predict connection existence of a simulated network from the associated distance and absolute density difference. Classification accuracy for predicting existence of connections in two species (macaque, blue; cat, green) by this classifier is shown. Accuracy was determined at each classification threshold (see Methods); here, we show mean accuracy across thresholds 0.750 to 0.975. Whether classification accuracy was better than chance was assessed by a permutation test, where classification accuracy was calculated for prediction from randomly permuted labels and a z-test was performed. A sign test was used to test whether the distribution of associated z-test p-values had a median value smaller than α = 0.05. The result of the sign test is indicated on top; black star: performance better than chance with median p < 0.05, red circle: performance not better than chance with median p >= 0.05. Box plots show distribution across 100 simulation runs per growth layout, indicating median (line), interquartile range (box), data range (whiskers) and outliers (crosses, outside of 2.7 standard deviations). See Table 4 for a summary. Abbreviations and background colours as in Table 1.

**Fig 9.**
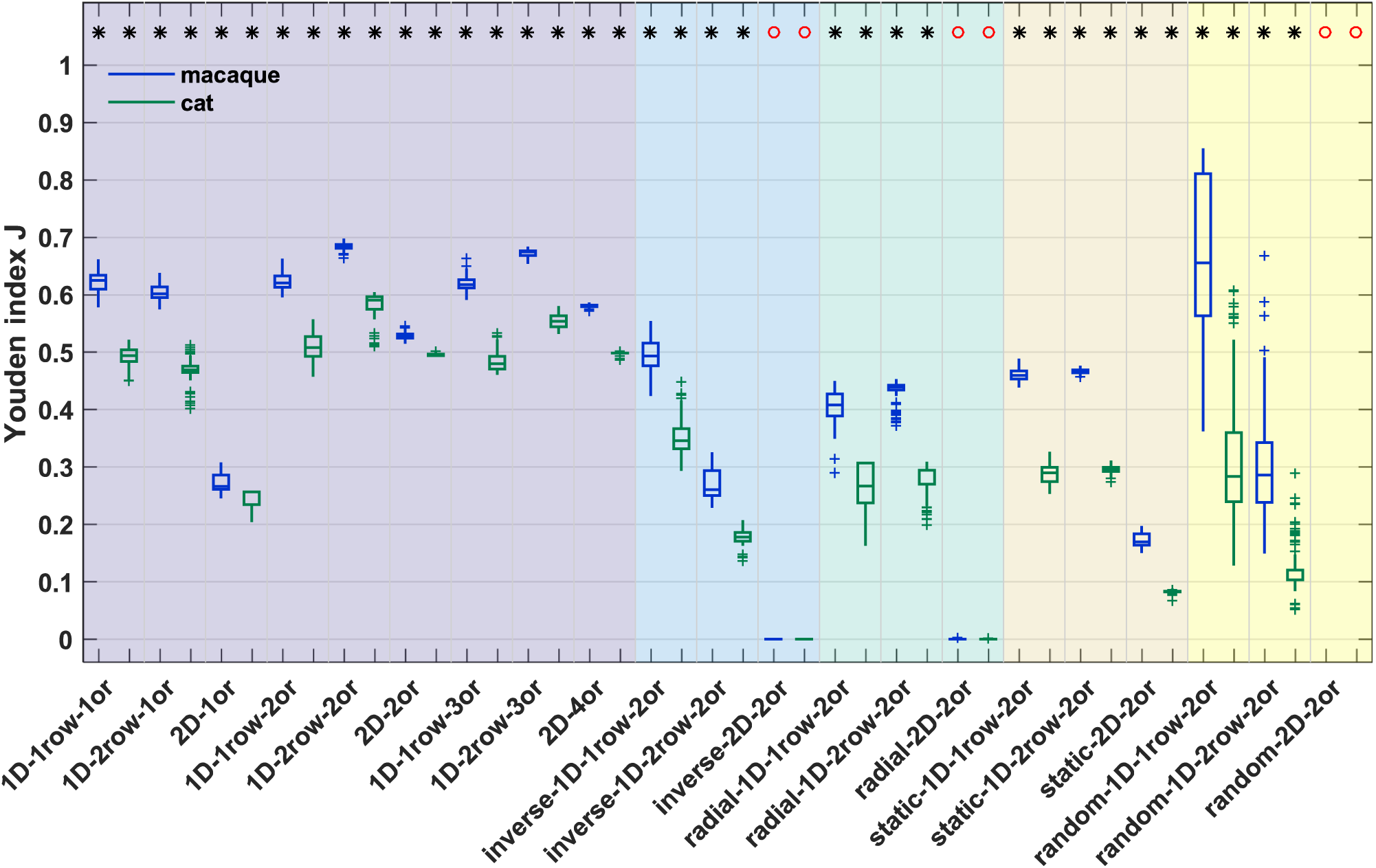
Youden index for prediction of empirical connection existence from simulation data. A classifier was trained to predict connection existence of a simulated network from the associated distance and absolute density difference. Youden index *J* for predicting existence of connections in two species (macaque, blue; cat, green) by this classifier is shown. Youden index *J* was determined at each classification threshold (see Methods); here, we show mean *J* across thresholds 0.750 to 0.975. Whether the Youden index was better than chance was assessed by a permutation test, where *J* was calculated for prediction from randomly permuted labels and a z-test was performed. A sign test was used to test whether the distribution of associated z-test p-values had a median value smaller than α = 0.05. The result of the sign test is indicated on top; black star: performance better than chance with median p < 0.05, red circle: performance not better than chance with median p >= 0.05. Box plots show distribution across 100 simulation runs per growth layout, indicating median (line), interquartile range (box), data range (whiskers) and outliers (crosses, outside of 2.7 standard deviations). See Table 4 for a summary. Abbreviations and background colours as in Table 1.

**Fig 10.**
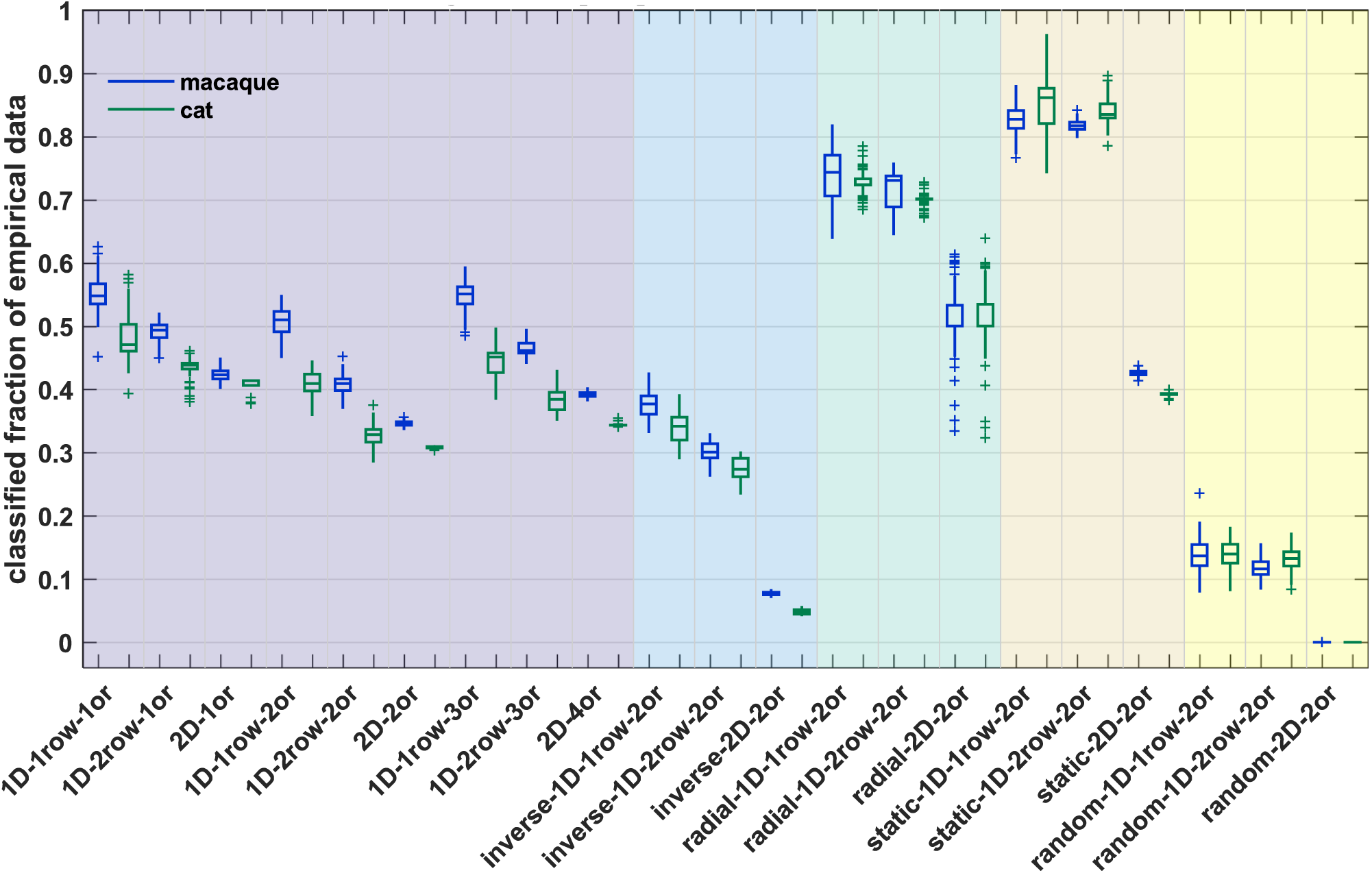
Percentage of empirical connectivity data that were classified from simulation data. A classifier was trained to predict connection existence of a simulated network from the associated distance and absolute density difference. This classifier was then used to predict connection existence in two species (macaque, blue; cat, green). Here, we show which fraction of the empirical data set was classified. This fraction differs across classification thresholds (see Methods); here, we show the mean fraction across thresholds 0.750 to 0.975. Box plots show distribution across 100 simulation runs per growth layout, indicating median (line), interquartile range (box), data range (whiskers) and outliers (crosses, outside of 2.7 standard deviations). See Table 4 for a summary. Abbreviations and background colours as in Table 1.

**Table 4.** Summary classification of empirical connectivity from simulated connectivity. This table lists the median values for classification accuracy, Youden index *J* and fraction of empirical connections classified as indicated by the box plots in Figures 8 to 10. For accuracy and Youden index, it additionally lists the associated median p-value of a z-test against chance performance as assessed by permutation analysis, as well as the z-value and p-value of a left-tailed sign test testing the distribution of z-test p-values for a median of α = 0.05. Background colours as in Table 1.

| growth layouts |  |  | Accuracy<br>(Fig. 8) |  |  |  |  |  |  |  | Youden index<br>(Fig. 9) |  |  |  |  |  |  |  | % classified<br>(Fig. 10) |  |
| --- | --- | --- | --- | --- | --- | --- | --- | --- | --- | --- | --- | --- | --- | --- | --- | --- | --- | --- | --- | --- |
|  |  |  | macaque |  |  |  | cat |  |  |  | macaque |  |  |  | cat |  |  |  | macaque | cat |
|  |  |  | prediction performance |  | validation of<br>'median p-value<br>against chance<br>performance': sign<br>test |  | prediction performance |  | validation of<br>'median p-value<br>against chance<br>performance': sign<br>test |  | prediction performance |  | validation of<br>'median p-value<br>against chance<br>performance': sign<br>test |  | prediction performance |  | validation of<br>'median p-value<br>against chance<br>performance': sign<br>test |  |  |  |
|  |  |  | median<br>accuracy | median p-value against<br>chance<br>performance | z-val | p-value | median<br>accuracy | median p-value against<br>chance<br>performance | z-val | p-value | median<br>*J* | median p-value against<br>chance<br>performance | z-val | p-value | median<br>*J* | median p-value against<br>chance<br>performance | z-val | p-value | median<br>fraction<br>classified | median<br>fraction<br>classified |
| realistically<br>oriented<br>gradient | 1D 1 row | 1 | 0.805 | 0.00E+00 | -9.90 | 2.08E-23 | 0.699 | 6.37E-18 | -9.90 | 2.08E-23 | 0.624 | 0.00E+00 | -9.90 | 2.08E-23 | 0.493 | 1.47E-17 | -9.90 | 2.08E-23 | 0.548 | 0.471 |
|  | 1D 2 rows | 1 | 0.781 | 4.54E-43 | -9.90 | 2.08E-23 | 0.640 | 6.45E-19 | -9.90 | 2.08E-23 | 0.601 | 9.04E-41 | -9.90 | 2.08E-23 | 0.468 | 7.02E-16 | -9.90 | 2.08E-23 | 0.493 | 0.438 |
|  | 2D | 1 | 0.811 | 3.14E-24 | -9.90 | 2.08E-23 | 0.830 | 2.57E-10 | -9.90 | 2.08E-23 | 0.265 | 1.95E-26 | -9.90 | 2.08E-23 | 0.256 | 8.99E-12 | -9.90 | 2.08E-23 | 0.423 | 0.414 |
|  | 1D 1 row | 2 | 0.854 | 0.00E+00 | -9.90 | 2.08E-23 | 0.820 | 6.03E-17 | -9.90 | 2.08E-23 | 0.620 | 0.00E+00 | -9.90 | 2.08E-23 | 0.507 | 3.90E-21 | -9.90 | 2.08E-23 | 0.510 | 0.409 |
|  | 1D 2 rows | 2 | 0.860 | 0.00E+00 | -9.90 | 2.08E-23 | 0.797 | 1.89E-16 | -9.90 | 2.08E-23 | 0.684 | 0.00E+00 | -9.90 | 2.08E-23 | 0.590 | 2.75E-19 | -9.90 | 2.08E-23 | 0.409 | 0.328 |
|  | 2D | 2 | 0.841 | 4.61E-35 | -9.90 | 2.08E-23 | 0.767 | 7.93E-15 | -9.90 | 2.08E-23 | 0.527 | 2.33E-41 | -9.90 | 2.08E-23 | 0.493 | 1.17E-17 | -9.90 | 2.08E-23 | 0.346 | 0.309 |
|  | 1D 1 row | 3 | 0.837 | 0.00E+00 | -9.90 | 2.08E-23 | 0.796 | 2.92E-17 | -9.90 | 2.08E-23 | 0.617 | 0.00E+00 | -9.90 | 2.08E-23 | 0.479 | 1.78E-20 | -9.90 | 2.08E-23 | 0.551 | 0.451 |
|  | 1D 2 rows | 3 | 0.843 | 0.00E+00 | -9.90 | 2.08E-23 | 0.757 | 4.00E-17 | -9.90 | 2.08E-23 | 0.673 | 0.00E+00 | -9.90 | 2.08E-23 | 0.553 | 6.09E-19 | -9.90 | 2.08E-23 | 0.461 | 0.384 |
|  | 2D | 3 | 0.847 | 5.33E-41 | -9.90 | 2.08E-23 | 0.767 | 6.32E-16 | -9.90 | 2.08E-23 | 0.579 | 0.00E+00 | -9.90 | 2.08E-23 | 0.497 | 8.88E-19 | -9.90 | 2.08E-23 | 0.392 | 0.343 |
| inverse<br>gradient | 1D 1 row | 2 | 0.782 | 7.70E-40 | -9.90 | 2.08E-23 | 0.586 | 3.59E-15 | -9.90 | 2.08E-23 | 0.492 | 7.06E-35 | -9.90 | 2.08E-23 | 0.345 | 1.31E-13 | -9.90 | 2.08E-23 | 0.376 | 0.341 |
|  | 1D 2 rows | 2 | 0.744 | 3.48E-35 | -9.90 | 2.08E-23 | 0.500 | 1.84E-13 | -9.90 | 2.08E-23 | 0.260 | 5.15E-22 | -9.90 | 2.08E-23 | 0.177 | 3.57E-08 | -9.90 | 2.08E-23 | 0.300 | 0.274 |
|  | 2D | 2 | 0.852 | 1.65E-22 | -9.90 | 2.08E-23 | 0.667 | 3.10E-06 | -9.90 | 2.08E-23 | 0.000 | -/- | -/- | 1.00E+00 | 0.000 | -/- | -/- | 1.00E+00 | 0.077 | 0.049 |
| radial | 1D 1 row | 2 | 0.739 | 7.89E-33 | -9.90 | 2.08E-23 | 0.671 | 4.24E-10 | -9.90 | 2.08E-23 | 0.407 | 6.28E-33 | -9.90 | 2.08E-23 | 0.266 | 2.32E-10 | -9.90 | 2.08E-23 | 0.743 | 0.723 |
|  | 1D 2 rows | 2 | 0.751 | 1.38E-34 | -9.90 | 2.08E-23 | 0.682 | 2.60E-11 | -9.90 | 2.08E-23 | 0.439 | 3.49E-35 | -9.90 | 2.08E-23 | 0.294 | 1.17E-11 | -9.90 | 2.08E-23 | 0.730 | 0.701 |
|  | 2D | 2 | 0.644 | 3.20E-01 | 5.50 | 1.00E+00 | 0.771 | 3.20E-01 | 5.90 | 1.00E+00 | 0.000 | 1.56E-02 | 0.00 | 5.00E-01 | 0.000 | 8.73E-03 | 0.00 | 5.00E-01 | 0.500 | 0.500 |
| static | 1D 1 row | 2 | 0.701 | 1.89E-39 | -9.90 | 2.08E-23 | 0.602 | 1.28E-11 | -9.90 | 2.08E-23 | 0.459 | 2.03E-40 | -9.90 | 2.08E-23 | 0.289 | 1.65E-11 | -9.90 | 2.08E-23 | 0.827 | 0.861 |
|  | 1D 2 rows | 2 | 0.703 | 1.12E-39 | -9.90 | 2.08E-23 | 0.602 | 6.91E-12 | -9.90 | 2.08E-23 | 0.465 | 8.41E-41 | -9.90 | 2.08E-23 | 0.294 | 1.24E-11 | -9.90 | 2.08E-23 | 0.817 | 0.834 |
|  | 2D | 2 | 0.869 | 1.40E-39 | -9.90 | 2.08E-23 | 0.897 | 6.17E-13 | -9.90 | 2.08E-23 | 0.169 | 1.86E-31 | -9.90 | 2.08E-23 | 0.083 | 4.35E-11 | -9.90 | 2.08E-23 | 0.426 | 0.393 |
| random | 1D 1 row | 2 | 0.880 | 1.70E-23 | -9.90 | 2.08E-23 | 0.722 | 4.65E-04 | -9.70 | 1.51E-22 | 0.655 | 1.29E-28 | -9.90 | 2.08E-23 | 0.283 | 1.43E-10 | -9.50 | 1.05E-21 | 0.137 | 0.139 |
|  | 1D 2 rows | 2 | 0.801 | 1.05E-19 | -9.90 | 2.08E-23 | 0.556 | 8.81E-03 | -9.90 | 2.08E-23 | 0.286 | 1.79E-16 | -9.90 | 2.08E-23 | 0.103 | 3.30E-10 | -9.70 | 1.51E-22 | 0.116 | 0.133 |
|  | 2D | 2 | 1.000 | 4.05E-02 | 0.71 | 7.60E-01 | 1.000 | 6.10E-02 | 3.88 | 1.00E+00 | -/- | -/- | -/- | 1.00E+00 | -/- | -/- | -/- | 1.00E+00 | 0.000 | 0.000 |

#### Accuracy

While classification accuracy is not a comprehensive measure to quantify classification performance, we included it to provide an overall impression of prediction quality. As seen in Figure 8 and Table 4, accuracy for most growth layouts surpassed chance performance, as assessed by a permutation analysis. Only the *random* and *radial 2D* growth layouts did not consistently reach better-than-chance accuracy. For classification of macaque connectivity, median accuracies that were better than chance ranged between 0.64 and 0.88, while the range of median accuracy for classification of cat connectivity was between 0.50 and 0.90. Comparing the different growth layouts, accuracy was generally higher for layouts with a *realistically oriented density gradient* than for *random, static, radial* or *inverse* growth layouts. The accuracies obtained for *realistically oriented gradient* growth layouts compared well to the accuracies we reported for the classification of empirical connectivity from the corresponding empirical structural measures, which were between 0.85 and 0.99 for the thresholds used here (cat, [34]; macaque, [36]). The better performance of *realistically oriented gradient* growth layouts was especially apparent if corresponding layouts were compared, for instance, in the macaque, the *random 1D 2rows* growth layout (median accuracy = 0.80) with the *realistically oriented density gradient 1D 2rows 2origins* growth layout (median accuracy = 0.86). Exceptions were, in the macaque, the *random 1D 1row* growth layout and the *inverse 2D* growth layout, as well as, in the macaque and in the cat, the *static 2D* growth layout, all of which had higher accuracy than the corresponding *realistically oriented* growth layout. However, all three growth layouts appeared inferior when their Youden index was considered (see below). Specifically, the *random 1D 1row* growth layout was very variable in terms of both accuracy and Youden index of classification performance, in contrast to the narrow distributions obtained for the *realistically oriented density gradient 1D 1row 2origins* growth layout. The *inverse 2D* growth layout reached a high accuracy for the prediction of macaque connectivity, but the Youden index showed that this did not lead to an overall prediction performance that was better than chance. Finally, for the prediction of both macaque and cat connectivity, the Youden index for the *static 2D* growth layout was below 0.2, indicating low overall prediction performance even though the obtained accuracies were very high.

#### Youden index

The Youden index, *J*, is a helpful summary measure of overall classification performance and affords a clear distinction between growth layouts. As seen in Figure 9 and Table 4, for most growth layouts *J* surpassed chance performance, as assessed by a permutation analysis. Exceptions here were the *random, radial* and *inverse 2D* growth layouts. Across the growth layouts with better-than-chance performance, classification performance ranged from poor to good, generally being somewhat higher for classification of the macaque connectome than for classification of the cat connectome. The highest values of *J* were reached for the layouts with growth along a *realistically oriented gradient*. In both species, performance for these growth layouts was moderate to good (macaque: median *J* = 0.53 − 0.68, cat: median *J* = 0.47 − 0.59). The only exception here was the *2D 1origin* growth layout, which reached only weak classification performance (macaque: median *J* = 0.27, cat: median *J* = 0.26). For the macaque, this performance compares well to the values of *J* that we previously reported for the classification of empirical connectivity from the corresponding empirical structural measures, which was 0.75 for the classification thresholds 0.85 through 1.00 [36]. Inclusion of the thresholds 0.75 and 0.80 would lower that value somewhat (cf. Fig. S2 in [36]).

Classification performance for the remaining growth layouts, namely the *random, static, radial* and *inverse* layouts, was low to moderate (median *J* macaque: generally < 0.49, median *J* cat: < 0.35). The difference to growth along a *realistically oriented gradient* was particularly apparent if corresponding layouts were compared. Growth layouts that reached moderate performance were the *static, radial* and *inverse 1D* growth layouts in the macaque. Their median *J* was still notably smaller than the median *J* value of the corresponding layout with growth along a *realistically oriented density gradient* (*1D 1row 2origins:* 0.62, *1D 2rows 2origins:* 0.68; *static 1D 1row:* 0.46, *static 1D 2rows:* 0.47; *radial 1D 1row:* 0.41, *radial 1D 2rows:* 0.44; *inverse 1D 1row:* 0.49, *inverse 1D 2rows:* 0.26; all values are for the macaque; cf. Table 4). The only exception to these observations was the *random 1D 1row* growth layout. In the macaque, this growth layout reached a median *J* of 0.65. However, the Youden index was also distributed very broadly, with a range of 0.36 to 0.85, indicating that classification performance was not consistently good, but volatile and strongly dependent on the particular instances of random neuron density patterns emerging in a given simulation.

**Classification performance varied with number of simulated growth origins**

To assess differences in classification performance in more detail, we compared the layouts with growth along a *realistically oriented gradient* among each other. Table 5 shows the results of a three-way analysis of variance for both accuracy and Youden index among the 9 growth layouts of set 1. We included the factors ‘species’ (macaque, cat), ‘growth mode’ (*1D 1row, 1D 2rows, 2D*) and ‘number of origins’ (1, 2, 3/4). For both accuracy and Youden index, the main effects of these three factors were significant. We performed post-hoc comparisons to describe the effect of the number of origins in more detail. As can be seen from Table 6, the comparisons showed that classification performance increased as the number of origins changed from one to two, but did not markedly increase further with addition of a third or fourth origin. In fact, for accuracy, there even was a slight decrease in the model estimate for three or four origins compared to two origins. This suggests that the network properties generated by growth around two origins were closer to empirical reality than those of networks grown around one origin, while a third or fourth origin did not further improve correspondence.

**Table 5.** Anova classification performance. A three-way analysis of variance was performed for both classification accuracy (see Fig. 8, Table 4) and Youden index *J* (see Fig. 9, Table 4), testing for effects of the factors ‘species’, ‘number of origins’, and ‘growth mode’. Sum Sq., Sum of squares; d.f., degrees of freedom; Mean Sq., mean squares = Sum.Sq. / d.f..

| Factor | Accuracy |  |  |  |  | Youden index *J* |  |  |  |  |
| --- | --- | --- | --- | --- | --- | --- | --- | --- | --- | --- |
|  | Sum Sq. | d.f. | Mean Sq. | F | Prob>F | Sum Sq. | d.f. | Mean Sq. | F | Prob>F |
| species | 2.00 | 1 | 2.00 | 1847.0 | 0 | 4.40 | 1 | 4.40 | 1283.7 | 0 |
| origins | 1.24 | 2 | 0.62 | 572.8 | 0 | 5.52 | 2 | 2.76 | 804.0 | 0 |
| growth mode | 0.28 | 2 | 0.14 | 128.7 | 0 | 8.03 | 2 | 4.02 | 1170.7 | 0 |
| Error | 1.94 | 1794 | 0.00 |  |  | 6.15 | 1794 | 0.00 |  |  |
| Total | 5.46 | 1799 |  |  |  | 24.11 | 1799 |  |  |  |

**Table 6.** Post-hoc comparisons classification performance. Post-hoc comparisons were computed to assess how classification accuracy and Youden index *J* were affected by the Anova factor ‘number of origins’. The upper section shows the marginal means estimated from the Anova-model. The lower section shows the results of the post-hoc tests for differences between the estimated means.

| model estimates | Accuracy |  | Youden index *J* |  |
| --- | --- | --- | --- | --- |
|  | estimated mean | standard error est. mean | estimated mean | standard error est. mean |
| 1 origin | 0.762 | 0.0013 | 0.450 | 0.0024 |
| 2 origins | 0.824 | 0.0013 | 0.568 | 0.0024 |
| 3/4 origins | 0.808 | 0.0013 | 0.567 | 0.0024 |

| post-hoc comparisons |  |  |  |  |
| --- | --- | --- | --- | --- |
|  | difference est. means | p-value difference est. | difference est. means | p-value difference est. |
| 1 vs 2 | -0.066 | 0 | -0.118 | 0 |
| 1 vs 3/4 | -0.046 | 0 | -0.117 | 0 |
| 2 vs 3/4 | 0.016 | 0 | 0.001 | 1 |

## Discussion

By performing comprehensive computational simulation experiments of how the network of interareal connections may develop during ontogenesis, we addressed the question of how cortico-cortical structural connections come to be closely related to the underlying substrate’s cytoarchitectural differentiation, an empirical observation made in multiple species [27, 30-37, 39-42]. The main component of our *in silico* model was a developing two-dimensional cortical sheet, gradually populated by neurons. To assess potential explanatory mechanisms, we varied the spatiotemporal trajectory of this simulated histogenesis. The rules governing axon outgrowth and connection formation, by contrast, were kept fixed across all variants of simulated histogenesis, so that the differences in outcome measures between spatiotemporal growth trajectories were introduced exclusively by the specifics of when and where neurons were generated.

To allow for straightforward interpretation of the simulation results, we applied network measures that were used in previous empirical studies, which allowed us to perform analyses on the simulated connectomes in the same way as we did on the empirical connectomes. Accordingly, the two characteristics of areas that were considered in the analyses of the final simulated network of interareal connections were their final position on the two-dimensional cortical sheet relative to other areas, measured as Euclidean distance, and their neuron density, which functioned as a surrogate for overall architectonic differentiation. Neuron density was expressed relative to the densities of other areas, that is, as density difference, for most analyses. We treated the existence of connections between areas as binary, thus, connections were considered as either absent or present.

### Spatiotemporal growth trajectories determine essential properties of the final connectome

We considered different spatiotemporal trajectories of how neurons populated the simulated cortical sheet. To recapitulate, simulated histogenesis proceeded according to five different sets of growth rules, with three to nine specific implementations per set (a total of 21 different growth layouts). These five sets were (1: *realistically oriented density gradient*) planar, expansive growth of the cortical sheet, with newer areas having successively higher neuron density; (2: *inverse gradient*) planar, expansive growth of the cortical sheet, with newer areas having successively lower neuron density; (3: *radial*) instead of planar growth, neurons started to populate all areas simultaneously and were added at a constant rate across the whole cortical sheet until each area reached its predetermined complement of neurons, with a final neuron density gradient identical to sets 1 and 4; (4: *static*) all neurons of the cortical sheet formed simultaneously, with a neuron density gradient identical to the final gradient of sets 1 and 3; (5: *random*) planar, expansive growth of the cortical sheet, with no ordered gradient of area neuron density around the two origins. To exclude effects specific to any particular implementation of these sets of growth rules, we considered three growth modes for each set: one-dimensional growth with one row of areas, one-dimensional growth with two rows of areas, and two-dimensional growth. For set 1, with a *realistically oriented density gradient*, we considered growth around one origin and three or four origins (for one-dimensional and two-dimensional growth modes, respectively) additionally to the growth around two origins that was used in all five sets.

These distinct spatiotemporal trajectories of cortical sheet growth led to considerable differences in the properties of the generated networks of structural connections. See Table 2 for an overall assessment of the results. While all growth layouts exhibited a clear decline in the relative frequency of present projections across larger distances, this measure correlated with density difference only for a subset of growth layouts (Fig. 5). Particularly, there was no consistent relationship for the *random*, *static* and *radial* growth layouts, while for oriented growth, both along a *realistically oriented density gradient* and along an *inverse gradient*, the relative frequency of present connections decreased with larger density differences between areas.

A more precise assessment of the extent to which distance and density difference determined connection existence was obtained by predicting simulated connectivity using binary logistic regression. Here, a similar picture as for relative connection frequency emerged from comparing McFadden’s Pseudo R^2^ values across growth layouts (Fig. 6). Distance was a better-than-chance predictor of connection existence for most growth layouts, as shown by the performance increase compared to a constant-only null model that is measured by McFadden’s Pseudo R^2^. In contrast, inclusion of density difference increased prediction performance only for the layouts with oriented growth (both along *realistically oriented* and *inverse density gradients*), but not for the *random, static* or *radial* growth layouts.

Finally, the growth layouts differed in whether neuron density correlated with area degree (Fig. 7). As before, for *random, static* and *radial* growth layouts, there was no consistent effect of neuron density on the measure of interest, in this case area degree. In contrast, there was a significant correlation with neuron density for layouts with oriented growth. This correlation was negative, as observed empirically, for growth layouts with a *realistically oriented density gradient*, but positive for growth layouts with an *inverse density gradient*.

In combination, these results demonstrate that the relation between neuron density, which is one crucial feature of the physical substrate in which connections are embedded, and cortico-cortical connections is strongly influenced by the precise spatiotemporal trajectory of cortex growth, which coincides with the time of connection formation. By manipulating where and when areas of varying neuron density were generated, we could observe a change in the extent to which connections of the simulated network were accounted for by the two factors of spatial proximity on the fully formed cortical sheet and the relative neuron density, indicating relative architectonic differentiation of areas.

### Realistic network properties emerge from empirically grounded growth trajectories

As described above, the extent to which spatial proximity and relative neuron density determined simulated connectivity strongly depended on the specific spatiotemporal trajectory of the simulated growth of the cortical sheet. Growth layouts that more closely mirrored the biological developmental trajectory of the mammalian cortical sheet led to closer correspondence of the simulation results with empirical observations on adult connectivity. This finding became especially apparent when we predicted empirical connectivity in two different mammalian species, cat and macaque, from regularities that were extracted from the simulated connectivity generated by the different growth layouts. Applying the regularities that emerged in our simulations to empirical data afforded a strong test of whether the simulations adequately captured ontogenetic processes and produced realistic networks. Our results showed that both of the aspects that were manipulated across growth layouts, the temporal trajectory of area growth as well as the direction of the neuron density gradient, were relevant for how well simulated connectivity allowed to predict empirical connectivity (Fig. 8, Fig. 9). First, it could be observed that growth layouts in which areas appeared successively around origins of neurogenesis (i.e., the *realistically oriented density gradient* growth layouts), were much better able to predict empirical connectivity than growth layouts with the same final neuron density gradient, but without the observed link between time of origin and neuron density (i.e., *static* and *radial* growth layouts). Second, in the presence of planar growth around origins, the direction of the neuron density gradient was crucial. This finding was indicated by the large decrease in prediction performance when comparing the *realistically oriented gradient* growth layouts with the *random* and *inverse density gradient* layouts. These sets of growth layouts both followed the same time course of cortical sheet expansion as the *realistically oriented gradient*, but with no relationship between time of origin and neuron density or a negative correlation between time of origin and neuron density, which contradicts the empirically observed positive correlation of time of origin with neuron density. Hence, the extent to which neuron density is well suited as a predictor of connectivity could be due to it reflecting neurodevelopmental time windows.

Third, our analyses revealed that the number of neurogenetic origins, around which new areas grew, influenced the correspondence to empirical connectivity (Table 5, Table 6). Growth around two origins arguably led to the best prediction performance: it was superior to growth around one origin for both accuracy and Youden index, and performed better than growth around three or four origins in terms of accuracy. For the Youden index, this performance difference was present, but too small to be meaningful or statistically significant. Thus, while correspondence between simulated and empirical connectivity clearly increased with the addition of a second origin of neurogenesis, there was at the very least no further performance increase with the addition of a third or fourth origin. Fourth, we observed that the overall level of prediction performance for the *realistically oriented density gradient* growth layouts was quite high, indicating that they afforded a good correspondence with empirical connectivity not only relative to the other growth layouts, but also in absolute terms. Therefore, a dual origin of neurogenesis and the resulting cytoarchitectonic gradients arguably are necessary components of the developmental mechanism for the empirically observed relations to hold (Fig. 11). These findings stress the importance of the theory of the dual origin of the cerebral cortex [38, 70] and the presence of multiple gradients of neurogenesis [56, 84], for the configuration of connectivity in the adult cortex.

Collectively, the presented results suggest that planar cortical sheet growth around two origins of neurogenesis and a systematic increase in neuron density with later time of origin are crucial determinants of the development of realistic cortico-cortical structural connections. Conversely, assuming that connection formation is a stochastic process with few constraints, as simulated here, the assumptions underlying the spatiotemporal growth trajectories of the *random, static, radial* and *inverse* growth layouts were shown not to mirror actual principles of cortical development.

**Fig 11.**
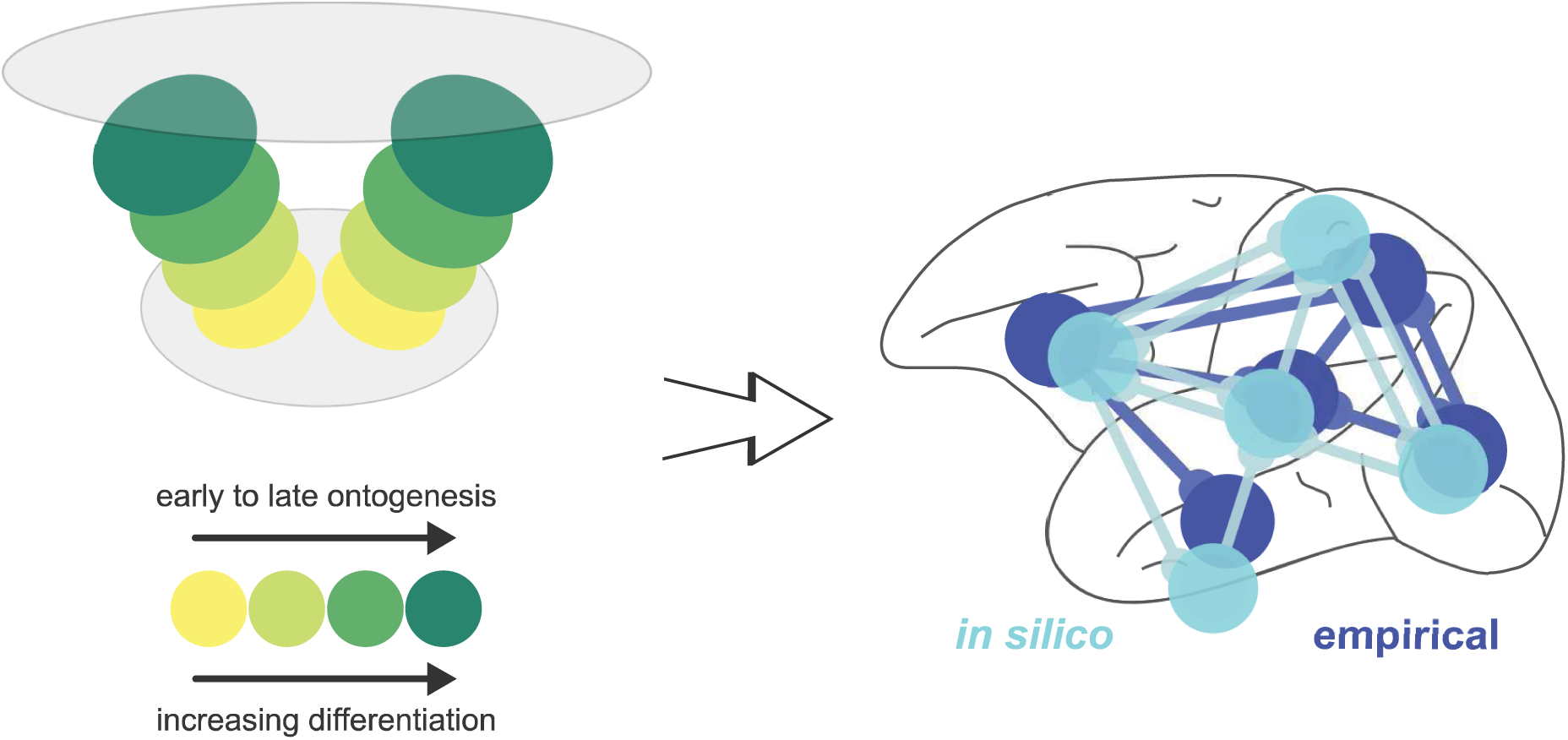
Number and relation of neurogenetic and architectonic gradients. A synthesis of all the results presented here indicates that the presence of two origins of neurogenesis, resulting in two neurogenetic (temporal) and architectonic gradients is necessary for the closer correspondence of the *in silico* model to the empirical relations between connectivity and architectonic differentiation. Importantly, the empirically observed relations are replicated *in silico* only when the less-to-more differentiated architectonic gradients align with early-to-late ontogenetic gradients. Hence, the suggested mechanism entails correspondence of neurogenesis and architectonic differentiation [36, 37, 40] and a dual origin of the cerebral cortex [38, 70].

### Simulation results validate the mechanistic explanations hypothesized to underlie the architectonic type principle

With the postulation of the architectonic type principle it was suggested that a close relationship between cortico-cortical connections and architectonic differentiation of the cortex might arise from the timing of neurogenesis [27], a process that occurs in close temporal proximity to the formation of connections. Specifically, it has been hypothesized that the relative time of generation of areas of different neuron densities affords them with different opportunities to connect with each other, thus imposing constraints on stochastically formed connections [29, 35]. This mechanism would be in line with findings in *Caenorhabditis elegans* [85] and rat cortex [46]. Moreover, a previous computational study already demonstrated that topological features, such as modular connectivity, may arise from the growth of connectivity within developmental time windows [86]. Thus, the main premise of this study, that spatiotemporal interactions in the forming cortex determine connectivity, has long been under consideration. Here, we provide the first systematic exploration of the possible mechanistic underpinnings of the ATP. We simulated multiple combinations of spatiotemporal growth trajectories of the cortical sheet and neuron density gradients, to probe from which of the combinations realistic connectivity emerged. Our results showed that, indeed, of the wide variety of examined spatiotemporal growth trajectories, the variant of the *in silico* model that led to the best correspondence with empirical observations was the one that was based on the same assumptions as the mechanism proposed to underlie the realization of the ATP. Hence, the underlying assumption that differences in neuronal density correspond to distinct time windows was not refuted in the model, and neuron density carried predictive power with respect to connectivity features only if such a relation between density and neurogenetic timing held. Our systematic simulation experiments, thus, distinctly corroborate the previously hypothesized mechanistic underpinnings of the ATP and contribute a conceptualization that can be scrutinized for similarities with, and distinctions from, actual ontogenetic processes. This approach opens up the possibility of characterizing in more detail how correlations between the structure of the cortex and cortical connections emerge, because all aspects of the process are observable. Further refinement of the simulation, for example by introducing species-specific histogenetic time courses, will enable the exploration of species differences or potentially the demonstration of invariance to changes in some aspects of ontogenesis. Another factor that could be probed is how robust the emergence of realistic connectivity is against changes in absolute neuron density, which varies considerably across species [65, 87]. From our simulations, it appears that temporal proximity of areas during neurogenesis underlies the positive relationship between similar neuron density and high connection probability. The close correlation between time of origin and architectonic differentiation described empirically (see Introduction) leads to a derivative correlation between temporal proximity of neurogenetic time windows and relative differentiation of cortical areas. Independent of this correlation, on a cortical sheet that expands around the origins of neurogenesis, areas with closer neurogenetic time windows tend to be spatially closer as well. Assuming connection formation is a stochastic process, which implies that connection probability declines with spatial distance, this process leads to a higher connection probability between areas that are generated during nearby time windows. Temporal proximity during neurogenesis would, thus, be the common antecedent determining both relative architectonic differentiation and connection probability, while those two factors would only be indirectly related. Temporal proximity, however, is difficult to measure, and it is, therefore, no surprise that the correlation between its two direct consequences has been empirically observed first. This chain of reasoning reveals how our modulation of the relationship between temporal proximity during neurogenesis and relative architectonic differentiation in the considered growth layouts could have caused the vastly different outcomes in connectivity described above.

In our simulations, we observed a relationship between the spatial proximity of areas and their likelihood to be connected, which appears to be an epiphenomenon of stochastic connection growth within a physically embedded system (c.f. [48, 88]). Distance is an inherent property of a spatially embedded system that cannot be removed from the implementation of spatial growth. However, in our simulation of cortical growth, the final distance between areas was not always an accurate measure of their distance during the time period of connection formation, which would be the factor that mattered principally for determining the likelihood by which two areas became connected. Since this distance during cortical sheet growth is correlated with areas’ final distance, there was also a correlation between final spatial proximity and connection probability. But this correlation does not genuinely describe the dependency of the stochastic growth process on distance, because inter-areal distance was not static, as implied by this measure. The distance measure relevant here, namely distance at the time of connection formation, would be challenging to measure empirically. Therefore relying on measures of final, adult distance and assuming a strong correlation between the two distance measures appears as a pragmatic strategy for empirical analyses.

### Simulating the development of laminar projection patterns

The present simulation experiments were designed to allow for the analysis of connection existence, that means, whether a possible connection between a pair of areas is simply present or absent in the final network. Naturally, axonal connections have many further properties beyond their simple existence; one prominent feature being the laminar distribution of the projection neurons’ somata and axon terminals in the areas of origin and termination, respectively. Laminar patterns of projection origin and termination are very well explained by the architectonic type principle (reviewed in [28, 29]), as has been demonstrated extensively in different species and cortical systems [27, 30-33, 35, 36, 39, 41, 42]. These conspicuous regularities most likely arise from fundamental developmental mechanisms, since they are ubiquitous and quite robust. This aspect becomes strikingly apparent in reeler mutant mice, where laminar connectivity patterns are largely correct [89-91]; shortly reviewed in [73], despite a systematic inversion (to ‘inside-out’) of neurons’ final laminar positions relative to the regular order that neurons typically assume according to their time of origin (‘outside-in’) [52, 89, 92-95]. However, the precise mechanisms through which laminar projection patterns become established are still under investigation. Further simulation experiments could, therefore, be helpful in evaluating potential candidate mechanisms. Expanding the simulation of cortical sheet growth to take into account the radial distribution of neurons across layers, as would be required for assessing laminar projection patterns, will afford the introduction of spatially and temporally more fine-grained features of neurogenesis and final architectonic differentiation. In addition to the planar gradient in neurogenetic time windows, which was taken into account in the present simulation experiments, this could mean to include the radial gradient in neurogenetic time windows that characterises neurons populating different layers [49, 50, 52, 61, 62]. Beyond the density of neurons in any given area, as considered here, structural variation could include a number of cellular morphological measures which have been shown to change systematically with overall density (e.g., [25]). Another feature that could prove to be relevant for the establishment of laminar projection patterns is the relative neuron density of cortical layers. As overall neuron density increases across the cortex, layer 2/3 becomes successively more pronounced [38, 96] and neuron density increases more in the supragranular layers than in the infragranular layers [66]. Thus, there is a shift in when, and in which layers, the majority of neurons is generated for areas of different overall density, which could affect laminar patterns, especially in interaction with the sequential growth of areas across the cortical sheet. Further, there is systematic variation in the size of pyramidal cell somata across the cortex, a phenomenon termed externopyramidization [70, 97]. Specifically, the ratio of soma size in supragranular to infragranular layers is much larger in areas of strong architectonic differentiation than in weakly differentiated areas (i.e., in weakly differentiated areas, infragranular pyramidal cells tend to be larger than supragranular cells, while the reverse is true for strongly differentiated areas). Hence, it seems that the laminar position of projection origins is aligned with relatively larger cell size in the candidate population for cortico-cortical connections, pyramidal cells. Since maintenance of long-distance connections between cortical areas is metabolically expensive [98, 99], relatively larger cell size conceivably is advantageous for their maintenance. Thus, as hypothesized before [43,100], externopyramidization might be linked to a shift in projection origins.

In addition to the above-mentioned properties of the cortical sheet itself, there are potential modifications of the stochastic formation of connections to be considered. First, the pruning of connections during later stages of development [101] was not taken into account in the present simulations. Laminar projection patterns may conceivably be affected by selective elimination of some axon branches but not others [102, 103]. Moreover, it has been observed that the time course of connection formation is not the same for all types of cells. Callosal projection neurons can reach their target areas without actually invading the gray matter, instead remaining in the white matter for a waiting period of days [104-106]. Similarly, waiting periods below the gray matter have been described for infragranular neurons projecting to area V4 from multiple areas in the ipisilateral hemisphere in macaques [107]. In contrast, supragranular neurons in the same tract-tracing experiments were found to invade the gray matter early, but many of them formed only transient projections that were subsequently eliminated. More generally, these and similar tract-tracing experiments have been interpreted to demonstrate different developmental profiles for axon outgrowth and connection formation in infra- and supragranular neurons [107-110]. In ‘feedback’ pathways, which according to the ATP can be conceptualised as projecting towards a relatively more differentiated area, extensive remodelling of laminar projection patterns until long after birth has been observed in a number of species (mouse, cat, macaque, human) and target areas [107-117]. This remodelling has been linked to activity-dependent maturation of pathways and the emergence of more refined perceptual capabilities [108, 117, 118]; e.g. reviewed in [119, 120]. This observation suggests that not all factors contributing to adult laminar projection patterns may be accessible in simulation experiments with time frames that are restricted to corticogenesis and initial axon outgrowth.

A further potential determining factor in the establishment of laminar projection patterns that warrants exploration is the possibility of genetic specification. The laminar position of projection targets might be regulated by genetically encoded factors. Numerous layer-specific transcription factors and neurotrophins have been described, which afford a precise targeting of specific layers or even cell types and cellular compartments (reviewed, e.g., in [121, 122]). Co-culture experiments using cortical explants have shown that appropriate laminar position of axon terminals was retained outside of the ontogenetic growth environment, that is, in the absence of regular temporal and spatial relationships. Accurate laminar specificity has been demonstrated, for example, for thalamo-cortical, geniculo-cortical, and cortico-spinal connections in co-culture (e.g., reviewed in [121]). Similarly, connections formed in co-culture of rat visual cortex explants were shown to conform to organotypic laminar distributions [123, 124]. Castellani and Bolz [125] elegantly demonstrated that organotypic and cell type specific projection patterns could be induced by membrane-associated factors through both induction and prevention of axon ingrowth and branching. Moreover, it has been shown that transcription factors can have population-specific effects, enlarging the range of potential interactions. For example, Castellani and colleagues [126] found that the membrane-bound protein Ephrin-A5 functioned as a repulsive axonal guidance signal in neurons destined to migrate to layer 2/3, while acting as a ‘branch-promoting’ signal in neurons destined for layer 6. These observations suggest that laminar patterns of projection terminations may not be entirely explicable by spatiotemporal interactions in the forming tissue, but are regulated by more prescriptive determinants.

### Limitations and future extensions

Our results illustrate how a mechanism linking the temporal order of neurogenesis across the cortex with the architectonic differentiation of areas could come to shape cortico-cortical connectivity such that it resembles the empirically observed connectivity features of mammalian connectomes. However, simulation experiments, as performed here, can only assess whether a suggested mechanism is principally feasible, and explore what its essential components might be. That is, such computational experiments put a candidate mechanism to the test and allow drawing some inferences about possible (and, importantly, impossible) ingredients, but they do not establish biological facts by themselves. Ultimately, only empirical observation of the ontogenesis of the cortex can establish how this developmental process unfolds. The possibility cannot be excluded that there may exist an unrelated mechanism working through features not considered here, which could cause the phenotype of interest, in our case the close relation between architectonic differentiation and connectivity. Generally, incorporating more empirical anchor points in a model gives the conclusions of a simulation study more significance. To triangulate a likely solution to the developmental puzzle of how axonal connections are organized, it is necessary to constrain potential mechanisms by as many observable features as possible. As discussed above, more processes that shape connectivity could be included in our *in silico* model of neural development, such as waiting periods for connection formation, a differential ability of cortical layers to retain connections (possibly linked to externopyramidization), the pruning of established connections, or the action of signalling molecules in attracting and repelling axons during connection formation. By integrating such processes, new insights could be gained into the emergence of further connection features such as laminar projection patterns and projection strengths.

We constrained our *in silico* model to represent a single cerebral hemisphere, hence our results only apply to ipsilateral, intra-hemispheric connections. Contralateral, inter-hemispheric axonal connections have also been reported to be well represented by the architectonic type principle [31, 37], although at generally lower connection strengths. The *in silico* model could be expanded by a second hemisphere which develops simultaneously. Since similar types of cortex in the two hemispheres would be formed at nearby points in time, but further apart in space, this setup would be expected to lead to the observed pattern of consistent, but weaker connectivity, if the principle holds that spatiotemporal interactions govern connectivity patterns.

We modelled the developing cortex as a two-dimensional sheet, across which axons grew until they met a target soma and formed a connection. In reality, the mammalian cortical sheet is not flat, but becomes at least curved, and often intricately folded, during corticogenesis. Moreover, axons are not positioned exclusively within the grey matter, but instead cover large distances through the white matter. These shortcuts between distant points on the cortical sheet imply that representing projection length as Euclidean distance between points on a flat cortical sheet is not accurate. Yet, regardless of how the concurrent processes of neurogenesis, axon formation and cortical folding affect each other [127, 128], measuring the precise lengths of projections in the adult cortex has so far not been straightforward. Hence, approximate measures have been employed, such as border distance on a cortical parcellation, Euclidean distance in three-dimensional space, or geodesic distance which accounts for some of projections’ confinement to white matter tracts. Euclidean distance on the simulated two-dimensional cortical sheet may, therefore, be a suitable surrogate measure for these approximate empirical measures. In line with this assumption, if cortical folding had a strong impact on our prediction of empirical data, it would be expected that performance in the less folded cat cortex would be better than in the more strongly folded macaque cortex. As this was not the case, we suspect that cortical folding and the resulting changes in projection lengths do not dramatically alter the spatiotemporal interactions which we hypothesize link architectonic differentiation and cortical connectivity. To further test this expectation, it would be interesting to predict connectivity data from a wider range of species, such as lyssencephalic rodents and humans, whose cortex is even more strongly folded than the macaque cortex.

Lastly, applying the classifier which was trained on simulated network data to predict empirical connectivity data resulted in better prediction performance for the macaque cortex than the cat cortex. Ultimately, there might be two reasons for this finding: Either the architectonic type principle characterises connectivity better in one of these species than the other, or the empirical measures that were used more faithfully capture the true structure in one of the species. Conceivably, adherence to the ATP might not be as pronounced in the smaller cat cortex, where both distances are shorter and therefore less distinctive, and there is less variation in total neuron number within the cortex due to a shortened neurogenetic interval [66]. Regarding the second possible reason, the structural measures from which we predicted connectivity were more detailed in the macaque cortex (neuron density and Euclidean distance) than in the cat cortex (structural type and border distance). Further experiments are therefore required to distinguish between these two explanations. Indeed, it would be intriguing to expand the prediction of empirical connectivity data from simulated networks to other species, preferably to mammals whose cortex is on either side of cat and macaque on the scales of size and degree of architectonic differentiation. Just as for assessing the impact of cortical folding, rodents and humans would be good candidates to identify the source of the observed difference in prediction performance.

## Conclusion

We performed simulations of cortical sheet growth and the concurrent formation of cortico-cortical connections, systematically varying the spatiotemporal trajectory of neurogenesis as well as the relation between architectonic differentiation and time of origin of neural populations. Our results showed that, for realistic assumptions about neurogenesis, successive tissue growth and stochastic connection formation interacted to produce realistic cortico-cortical connectivity. This finding illustrated the fact that precise targeting of interareal connection terminations was not necessary for obtaining a realistic replication of connection existence within a cortical hemisphere. Instead, spatiotemporal interactions within the structural substrate were sufficient if a small number of empirically well-grounded assumptions were met, namely (i) planar, expansive growth of the cortical sheet as neurogenesis progressed, (ii) stronger architectonic differentiation for later neurogenetic time windows, and (iii) stochastic connection formation. We, thus, demonstrated a possible mechanism of how relative architectonic differentiation and connectivity become linked during development. These findings support hypotheses advanced in previous reports about the mechanistic underpinnings of the architectonic type principle [27, 29, 35, 40]. The successful prediction of connectivity in two species, cat and macaque, from our simulated cortico-cortical connection networks further underscores the generality of the ATP and the wide applicability of its explanation of connectivity in terms of relative architectonic differentiation.

## Methods

We first describe the variants of the *in silico* model we considered and how we simulated the formation of cortico-cortical connections on a forming cortical sheet, representing a single hemisphere. We then detail how we analysed the resulting simulated networks.

Connection formation was simulated to take place on a two-dimensional, rectangular cortical sheet, where neuron somata and axon terminals were assigned two-dimensional coordinates without spatial extent. Somata were arranged in rectangular cortical areas which differed in their surface density of neurons. Neuron density has been shown to be a good indicator of a cortical area’s overall degree of architectonic differentiation [40] and has been used previously to relate differentiation to connectivity in the macaque brain (e.g. [35, 36]). Hence, we used neuron density as a central marker for architectonic differentiation, with larger neuron density corresponding to a stronger degree of differentiation. We did not adjust the absolute magnitude of neuron density to empirical values, but did choose the range of neuron densities such that it was similar to empirically observed variation in neuron densities across the cortex, with about a five-fold increase between areas of lowest and highest neuron density (cf. [35]). We implemented neuron density as number of somata per unit area of cortical sheet (#/arbitrary unit^2^). All cortical areas were defined to be of the same size. From these two constraints on neuron density and area size, it followed that areas of different densities contained different numbers of neurons. Within an area, somata were spaced equidistantly.

### Variants of the *in silico* model

The generation of the cortical sheet across time was simulated in a number of different settings of the *in silico* model, here called variants or growth layouts. These growth layouts systematically differed in where and when neurons were generated on the forming cortical sheet, that is, they had different spatiotemporal growth trajectories. Below, we describe all growth layouts and their correspondence to neurodevelopmental findings in detail. An overview is provided in Table 1, and Figure 2 as well as Supplementary Figure S1 give a visualisation of cortical sheet development over time for the different growth layouts.

All considered spatiotemporal growth trajectories were grouped into five sets of growth layouts. These sets differed with respect to whether cortical areas were generated by planar, expansive growth, whether there was radial growth, and in the final gradient of neuron density around neurogenetic origins.

In growth layouts with planar growth, the cortical sheet expanded, as, with each growth event, new cortical areas emerged around neurogenetic origins. Each new cortical area was grown within one time step, thus all constituent neurons appeared on the cortical sheet simultaneously. Neurogenesis occurred on the outer fringes of the portion of the cortical sheet already generated by each origin of neurogenesis. For more than one neurogenetic origin, this process entailed that newly generated areas moved previously generated areas apart on the cortical sheet, increasing the spatial distance in between them. Thus, planar growth mimicked the empirically observed planar gradient in onset of neurogenesis (see Introduction).

Radial growth, in contrast, did not expand the cortical sheet over time. Here, the cortical sheet had its final dimension already at the start of corticogenesis and cortical areas did not differ with respect to the time of onset of neurogenesis, but instead in the length of their neurogenetic interval. During each growth event, neurons were added at a constant rate across the entire cortical sheet. Areas with lower neuron density finished generating their complement of neurons earlier in time than areas with a higher neuron density, which needed to generate a larger number of neurons. Radial growth thus reproduced an alternative interpretation of studies of neurogenetic timing (see Introduction).

Growth events, during which the cortical sheet was generated, were distributed across the fixed simulated length of time. For both planar and radial growth, they were timed in such a manner that all neurons had grown after one third of the simulation length, and the remaining time steps could be used for connection formation by all neurons.

These three main properties of spatiotemporal growth of the cortical sheet were combined in the five sets of growth layouts, with each set containing three (or in one case nine) growth layouts, as follows: The first set, the *realistically oriented density gradient* growth layouts, grew by planar growth. Here, newly generated areas were of higher neuron density than previously grown areas. That is, there was a positive correlation between time of origin and neuron density, which appeared as a distinct gradient in neuron density around the neurogenetic origins on the final cortical sheet. The second set, the *inverse neuron density gradient* growth layouts, grew by planar growth like sets 1 and 5. However, in these *inverse gradient* growth layouts, newly generated areas were of lower neuron density than previously grown areas, that is, there was a negative correlation between time of origin and neuron density. The third set, the *radial* growth layouts, grew by radial growth. The final density gradient was identical to sets 1 and 4, but for the *radial* growth layouts, this pattern was caused by a positive correlation between length of the neurogenetic interval and neuron density, instead of a correlation between the time of onset of neurogenesis and neuron density. The fourth set, *static* growth layouts, did not in fact grow at all. All neurons were grown during the first growth event, thus the cortical sheet was fully formed from the beginning of the simulation. The final density gradient was identical to sets 1 and 3. Finally, in the fifth set, the *random* growth layouts, the cortical sheet grew by planar growth. The resulting final cortical sheet had no directed gradient of neuron density around the neurogenetic origins. Instead, each newly generated area was randomly assigned a neuron density. Possible density values were drawn from the neuron densities found on the final cortical sheet of the first set, *realistically oriented neuron density gradient*.

For each of these five sets, we implemented three different growth modes to mitigate influences of any specific choice of spatial implementation. Each growth mode was implemented around two neurogenetic origins. The three growth modes were as follows: First, one-dimensional growth with one row of areas (*1D 1row* growth layouts), where new areas grew to the left and right of neurogenetic origins (i.e., along the x-dimension of the cortical sheet) and there was only one row of cortical areas. Second, we implemented one-dimensional growth with two rows of areas (*1D 2rows* growth layouts), where, again, areas were added to the left and right of neurogenetic origins, but there were two rows of areas stacked in the y-dimension of the cortical sheet. Third, we implemented two-dimensional growth (*2D* growth layouts), where new areas were added on all sides of neurogenetic origins (i.e., in both the x- and y-direction of the cortical sheet). In this growth mode, each successive growth event led to an exponentially increasing number of added areas, and for set 1, *realistically oriented density gradient*, an unproportionally high number of areas of the highest neuron density, which did not accurately reflect the composition of the mammalian cerebral cortex. However, as stated above, we simulated the different growth modes to alleviate side-effects that might unintentionally arise from any particular spatial layout. Considering results across these specific implementations vastly reduced the risk of misinterpretation. We therefore included the two-dimensional growth mode despite its unrealistic rendering of the cortical sheet as a further control.

As mentioned before, each of the 15 growth layouts that were described so far was implemented around two origins of neurogenesis (5 sets × 3 growth modes × 1 number of origins). For set 1, *realistically oriented neuron density gradient*, we additionally considered two different numbers of origins for each growth mode. Specifically, we included growth around one neurogenetic origin and growth around three or four neurogenetic origins for *1D* and *2D* growth modes, respectively. These further six growth layouts allowed us to test whether the exact number of neurogenetic origins meaningfully influenced final connectivity.

Thus, we considered a total of 21 growth layouts (5 sets × 3 growth modes × 1 number of origins + 1 set × 3 growth modes × 2 numbers of origins). We simulated 100 instances of the spatiotemporal development of each of these 21 growth layouts.

#### Correspondence to empirical observations

The five sets were designed to correspond to some aspects of empirical neurodevelopmental findings and to violate others. Set 1, which features planar growth and a *realistically oriented density gradient*, represents a fiducial reproduction of the empirically grounded assumptions we described in the Introduction and thus mimics the mechanistic underpinnings that were previously hypothesised for the architectonic type principle [27, 29, 35, 40]. The other four sets deviate from this most realistic set in different ways. Sets 2 and 5, with *inverse* and *random* density gradients, respectively, test how the specifics of the neuron density gradient affect connectivity in the presence of planar growth. Set 4, the *static* growth layouts, examine how the absence of planar growth affects connectivity if the neuron density distribution remains unchanged. Set 3, *radial* growth layouts, contrast planar growth with radial growth, while the final neuron density distribution again remains unchanged.

### Connection formation

Axons randomly grew across the cortical sheet and stochastically formed synaptic connections (similar to, e.g., [48]; also see [45]). Each neuron was assigned one axon terminal, which was initially located at the respective soma position. With each time step of the simulation, the axon extended by a fixed length at a random angle, and the position of the axon terminal changed accordingly. Once axon terminals left the cortical area their parent soma was located in, they were free to form a synapse with any neuron soma they encountered. Since both terminals and somata were defined by point-coordinates, a synapse was formed once the axon terminal approached a soma closer than a defined maximal distance. Upon synaptic contact, an axon stopped growing and the now occupied axon terminal remained at the location of the contacted soma for the remainder of time steps. To further increase stochasticity, we imposed a connection probability of 90% on potential synaptic contacts. Thus, in 90% of cases, a synapse successfully formed once the terminal was close enough to a soma, but in a randomly chosen 10% of cases, no synapse formed at this time step and the axon continued to grow. If soma positions changed because the cortical sheet grew, axon terminals (both occupied and unoccupied) were shifted with the cortical area they found themselves in at the time, and synaptic contacts were retained. This procedure of axon growth and synapse formation was not modified across variants of the *in silico* model.

Different parameters of the axon growth process interacted to determine how fast axon terminals made synaptic contacts. This included for example the increase in axon length per time step and the maximal distance for synapse formation. In pilot runs of the simulation, we calibrated the relevant parameters such that after the fixed simulated length of time, most axon terminals (>99.9%) had made synaptic contact and final interareal connectivity fell into a range comparable to empirical reports [22, 34, 83]. This calibration resulted in slightly different parameter values for *1D* and *2D* growth modes, but the same values were used in all simulation instances within these growth modes.

### Features of the simulated cortical sheet

From the final state of the simulated cortical sheet, we extracted a number of features that were analogous to measures used in previous analyses of the mammalian cortex.

First, we collapsed the axonal connections between individual neurons into a simulated connectome, which contained information about the existence of all possible area-wise connections. Thus, we constructed a complete binary connectivity matrix where connections were entered as either absent or present.

Second, we extracted the two relevant structural measures from the final cortical sheet. The first measure was each area’s neuron density, and derived from that the difference in neuron density between area pairs, where density difference = density_area of origin_ - density_area of termination_. For most analyses, we considered the undirected equivalent, the absolute value of density difference, which indicates the magnitude of the difference in neuron density between two areas. These two measures were equivalent to measures of architectonic differentiation previously used in studies examining mammalian cortical connectivity, such as neuron density difference (e.g., [6635 the log-ratio of neuron densities [36] or difference in cortical type, which is an ordinal measure of architectonic differentiation (e.g., [33-35]). The second measure was the spatial proximity between pairs of areas, which we calculated as the Euclidean distance between areas’ centres of mass. This measure was equivalent to measures of spatial proximity we used in previous empirical studies (e.g., [35, 36]). Since distance is an undirected measure, each analysis that included distance required the use of the undirected measure of neuron density difference, its absolute value.

### Analyses

We performed the described analyses for each of the 100 instances that were simulated for each growth layout and aggregated results across instances. For the simulations and analyses we used Matlab R2016a (The MathWorks, Inc., Natick, MA, USA).

#### Relative frequency of present connections

To gain an overview of how present and absent connections were distributed across the range of possible absolute density differences and distances, we computed the relative frequency of present connections. To do this, we divided the range of each structural measure in up to 10 bins and computed the fraction of present connections in each bin as relative frequency = number of present connections / (number of present connections + number of absent connections). For distance, we always used 10 bins. For absolute density difference, we used 10 bins where possible, but we had to chose a lower number of bins if the particular growth layout had been implemented with a small number of area density tiers. This was for example the case in the *2D 4origins* growth layout, where the exponential increase in the number of areas with each growth event caused us to restrict the simulation to four growth events, and thus four different levels of neuron density. To assess whether there was a systematic relation between the relative frequency of present connections and the respective structural measure, we then computed Spearman rank correlations of the computed fractions across all bins. We show the resulting distribution of correlation coefficients ρ and report median ρ- and p-values averaged across simulation instances. To determine whether the rank correlation was consistently significant across instances, we computed a left-tailed sign test for each growth layout. Specifically, we tested whether the group of 100 p-values obtained from the rank correlations for each instance had a median value smaller than a significance threshold, α_Spearman_ = 0.05. We considered the sign test significant below α_sign_ = 0.05, and in these cases rejected the null-hypothesis that the median of the group of p-values was not smaller than α_Spearman_. For the sign test, we report the test statistic z and the corresponding p-value.

#### Prediction of simulated connectivity data

To assess how well density difference and distance accounted for the simulated interareal connectivity, we used binary logistic regression, a classification algorithm for distinction between two classes of cases. That is, we endeavored to predict the existence of simulated connections from the structural properties of the corresponding simulated cortical sheet. We considered four combinations of predicting factors: First, a null model which included only a constant and amounted to chance performance. Second and third, we further included either absolute density difference or distance as predicting factors. Thus, we constructed two models with two predicting factors each, testing the effect of each individual structural measure on classification performance. In a fourth model, we included all three predicting factors, that is, a constant and both structural measures, testing their joint classification performance. Prior to inclusion, both structural measures were transformed to z-scores, that is, we subtracted the respective mean and then divided by the respective standard deviation. To evaluate how much each predicting factor contributed to classification performance, we computed McFadden’s Pseudo R^2^ = log-likelihood_model_ / log-likelihood_null model_. The log-likelihood for each model captures how well its predictions correspond to the actual data, with larger values indicating a better correspondence. McFadden’s Pseudo R^2^ thus indicates how much better prediction performance becomes with the inclusion of further predicting factors, relative to chance performance. Values of McFadden’s Pseudo R^2^ of 0.10 and above were considered a moderate increase in prediction performance, values of 0.15 and above were considered adequate, and values from 0.20 on were considered a very high increase in prediction performance [129].

#### Area degree

We assessed one topological property of areas, their degree, which has previously been reported to be related to architectonic differentiation [34, 36]. Area degree indicates how many connections are maintained by an area, and we computed it as the sum of afferent and efferent connections for each area. Since degree is not a relational property and hence applies to a single area and not a pair of areas, we related it to neuron density but not to spatial proximity. Analogous to our previous analyses, we computed a Spearman rank correlation between area degree and neuron density to assess whether there was a relation between the two. We show the resulting distribution of correlation coefficients ρ and report median ρ- and p-values averaged across simulation instances. To determine whether the rank correlation was consistently significant across instances, we computed a left-tailed sign test for each growth layout, as described above for relative connection frequencies. The same significance thresholds applied here.

#### Prediction of empirical connectivity data

To assess how well the relationships between simulated connectivity and simulated structural measures translated to empirically observed relations in the mammalian cortex, we used classifiers trained on the simulated data to predict empirical connectivity data. To this end, we used two data sets of ipsilateral corticocortical connectivity (i.e., connections within a hemisphere), analyses of which we have published previously. These were the most extensive and up-to-date connectivity data sets available for the macaque [83] and cat cortex [130], acquired using retrograde tract-tracing experiments. Here, we considered these connectivity data as a binary measure of connection existence. For both data sets, measures of architectonic differentiation and spatial proximity were available. Briefly, in the macaque, we used the absolute log-ratio of neuron density and Euclidean distance between areas as the equivalents of the absolute density difference and Euclidean distance obtained from the simulations and included 1128 empirical data points in our analyses. In the cat, these measures were represented by the absolute difference in cortical type, an ordinal ranking of areas by architectonic differentiation, and the border distance between areas, which quantifies the shortest distance between two areas based on a given parcellation of the cortex. Here, we included 954 empirical data points in our analyses. See our previous reports for a detailed description of the connectivity data as well as the structural measures [34, 36]. To be able to apply the two simulated structural measures to the empirical measures despite their different scales, we transformed all three pairs (simulated, macaque, cat) to z-scores by subtracting the respective mean and then dividing by the respective standard deviation.

For each instance of each growth layout, we trained a classifier to predict simulated connection existence from the z-scores of simulated relative architectonic differentiation (i.e., absolute density difference) and spatial proximity (i.e., distance), using a support vector machine with a linear kernel function and the assumption of uniform prior probabilities for the two learned classes. We then applied the trained classifier to the z-scores of empirical relative architectonic differentiation (i.e., absolute log-ratio of neuron density and absolute type difference, respectively) and spatial proximity (i.e., Euclidean distance and border distance, respectively), separately for the macaque and the cat, and obtained posterior probabilities that a connection was present, p_present_. We then used two classification rules, derived from a common threshold probability p_threshold,_ to label empirical data points as either absent or present. We assigned the status ‘present’ to all empirical connections whose posterior probability exceeded the threshold probability, that is, data points with p_presen_ > p_threshold_. Alternatively, we assigned the status ‘absent’ to all empirical connections whose posterior probability was sufficiently low, that is, data points with p_present_ < 1- p_threshold_. These two rules excluded a range of posterior probabilities where classification was not confident enough to warrant a prediction, which entailed that not all empirical connections were assigned a predicted label for each simulation instance. Additionally to the measures that we used to quantify prediction performance, we therefore report the fraction of available empirical data points that were actually classified. To mitigate influences of any one threshold probability, we considered ten threshold probabilities, increasing in step sizes of 0.025 from p_threshold_ = 0.750 to p_threshold_ = 0.975, and report results averaged across thresholds for each simulation instance.

We assessed prediction performance through two measures, accuracy and the Youden index, *J*. We calculated these measures at each threshold and report results averaged across all ten thresholds. Accuracy was computed as the fraction of predictions that were correct, that is, accuracy = number of correct predictions / (number of correct predictions + number of incorrect predictions). The Youden index *J* [131, 132] is a more comprehensive summary measure which takes into account both sensitivity (true positive rate) and specificity (true negative rate), with *J* = sensitivity + specificity − 1. The Youden index is a measure of how well a binary classifier operates above chance level, where *J* = 0 indicates chance performance and *J* = 1 indicates perfect classification. Below values of 0.25, the Youden index was considered to indicate negligible classification performance, values of 0.25 and above were considered weak performance, values of 0.40 and above were considered moderate performance, and values of *J* above 0.50 were considered to indicate good classification performance.

We show the distribution of resulting mean values of accuracy and Youden index across the ten thresholds, and report the median values of these distributions across the 100 instances for each growth layout. In the following, we describe the procedure that we followed to validate the two classification performance measures, assessing how they compared against chance performance. An overview is provided in Figure 3. Within each simulation instance, we performed a permutation analysis at each threshold to determine how the accuracy or Youden index at this threshold compared to chance performance. To this end, we randomly permuted the labels of the empirical data points, so that there was no association any more between the predictive variables and connection existence, and then applied the classification procedure again, computing accuracy and Youden index to quantify chance performance. We repeated this for 100 permutations of the data labels, so that, for both measures, we obtained a distribution of values that represented chance performance at each threshold. To test whether the corresponding classification performance measure was likely to be from this distribution, we first fit the chance performance distribution to a normal distribution, obtaining an inferred mean value and standard deviation. We then performed a two-tailed z-test, which tests whether a particular value comes from a population with a particular mean, which in this case was the fitted distribution of performance measures obtained from the permutation analysis. If the test was significant at α_z-test_ = 0.05, we rejected the null hypothesis that the actual performance measure at the given threshold came from the fitted distribution of chance performance. Since the z-statistic was never smaller than 0 if the p-value was below α_z-test_, we then inferred that the actual performance was better than chance performance at a given threshold. We then averaged the p-values obtained from the z-tests across thresholds by computing their median. Thus, for each growth layout, we obtained distributions of 100 (one per instance) mean performance measures and as many associated median p-values validating them against chance performance.

To determine whether these median p-values were consistently significant across instances, we computed a left-tailed sign test for each growth layout. Specifically, we tested whether the group of 100 median p-values obtained from the z-tests at each threshold for each instance had a median value smaller than α_z-test_. We considered the sign test significant at α_sign_ = 0.05, and in these cases rejected the null-hypothesis that the median of the group of p-values was not smaller than α_z-test_. For the sign test, we report the test statistic z and the corresponding p-value for each growth layout.

Finally, to assess how the two classification performance measures accuracy and Youden index were affected by the number of origins independent of growth mode and the considered species, we computed a three-way analysis of variance on the performance measures from growth layouts with a realistically oriented density gradient (which were the only ones where number of origins ever differed from two). We included three factors: ‘species’, with the levels macaque and cat; ‘growth mode’, with the levels *1D 1row, 1D 2rows* and *2D;* and ‘number of origins’, with the levels 1, 2 and 3 or 4 (for *1D* and *2D* growth modes, respectively). We report the F-statistic and associated p-value for each factor and considered a main effect significant at α_ANOVA_ = 0.05. To examine the main effect of ‘number of origins’ in more detail, we estimated marginal mean values from the analysis of variance model. These reflect a model estimate of the mean value for each level of ‘number of origins’ across all levels of the remaining factors. We subsequently performed post-hoc comparisons between these model estimates of marginal mean values, which revealed specific differences between levels. The post-hoc comparisons were Bonferroni corrected for multiple tests and considered significant at an adjusted threshold of α_post-hoc_ = 0.05/3 = 0.0167.

## Supplementary Figure Captions

**Supplementary Figure S1.** Developmental trajectories of all 21 growth layouts. Illustration of the spatiotemporal growth trajectory for each growth layout. The successive population of the cortical sheet with neurons is shown for the first three growth events. For static growth, all neurons grow simultaneously, hence only one growth event is shown.

**Supplementary Figure S2.** Correlation of relative connection frequency with distance and absolute density difference for all growth layouts. Distribution of absent and present connections across distance (left panels) and absolute density difference (right panels) for all growth layouts. Absolute numbers of absent and present projections (bars) are depicted alongside the corresponding relative frequency of present connections (diamonds). Simulation instances were chosen to be representative of the median values shown in Figure 5. Spearman rank correlation results for each particular instance are shown on top of each plot. A.u.: arbitrary unit. Abbreviations and background colours as in Table 1.

**Supplementary Figure S3.** Correlation area degree with neuron density for all growth layouts. Variation of area degree (number of connections) across areas’ neuron density is shown. Simulation instances were chosen to be representative of the median values shown in Figure 7. Spearman rank correlation results for each particular instance are shown on top of each plot. A.u.: arbitrary unit. Abbreviations and background colours as in Table 1.

